# Parallel Expansion and Divergence of an Adhesin Family in Pathogenic Yeasts Including *Candida auris*

**DOI:** 10.1101/2022.02.09.479577

**Authors:** Rachel A. Smoak, Lindsey F. Snyder, Jan S. Fassler, Bin Z. He

## Abstract

Opportunistic yeast pathogens evolved multiple times in the Saccharomycetes class, including the recently emerged, multidrug-resistant *Candida auris*. We show that homologs of a known yeast adhesin family in *Candida albicans*, the Hyr/Iff-like (Hil) family, are enriched in distinct clades of *Candida* species as a result of multiple, independent expansions. Following gene duplication, the tandem repeat-rich region in these proteins diverged extremely rapidly and generated large variations in length and β-aggregation potential, both of which were known to directly affect adhesion. The conserved N-terminal effector domain was predicted to adopt a β-helical fold followed by an α-crystallin domain, making it structurally similar to a group of unrelated bacterial adhesins. Nonsynonymous-to-synonymous substitution rate analysis of the effector domain in *C. auris* revealed relaxed selective constraint and signatures of positive selection, suggesting functional diversification after gene duplication. Lastly, we found the Hil family genes to be enriched at chromosomal ends, which likely contributed to their expansion via ectopic recombination and break-induced replication. We hypothesize that the expansion and diversification of adhesin families are a key step toward the emergence of fungal pathogens and also generate variation in adhesion and virulence within and between species.

## Introduction

*Candida auris*, a newly emerged multidrug-resistant yeast pathogen, is associated with a high mortality rate (up to 60% in a multi-continent meta-analysis (Lockhart et al. 2017)) and has caused multiple outbreaks across the world (CDC global *C. auris* cases count, February 15th, 2021). As a result, it became the first fungal pathogen to be designated by CDC as an urgent threat (CDC 2019). The emergence of *C. auris* as a pathogen is part of a bigger evolutionary puzzle: *Candida* is a polyphyletic genus that contains most of the human yeast pathogens. Phylogenetically, species like *C. albicans, C. auris* and *C. glabrata* belong to distinct clades with close relatives that either don’t or only rarely infect humans (Fig. 1A). This strongly suggests that the ability to infect humans evolved multiple times in yeasts (Gabaldón et al. 2016). Because many of the newly emerged *Candida* pathogens are either resistant or can quickly evolve resistance to antifungal drugs (Lamoth et al. 2018; Srivastava et al. 2018), it is urgent to understand how yeast pathogenesis arose and what increases their survival in the host. We reason that any shared genetic changes among independently derived *Candida* pathogens will reveal key factors for host adaptation.

**Figure 1.**
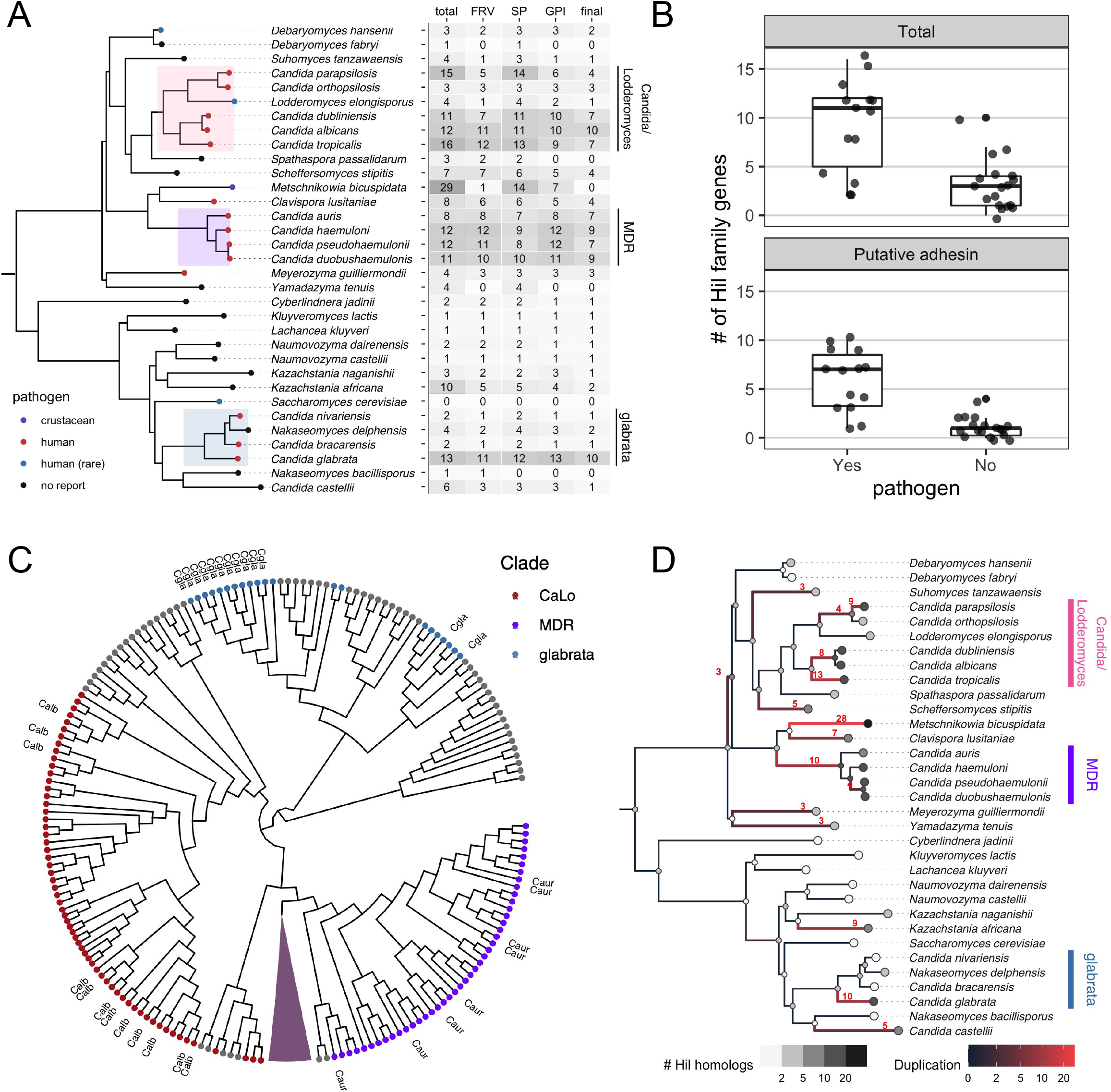
Phylogenetic distribution of the yeast Hil family and its parallel expansion in independently derived pathogenic *Candida* species. A) Species tree is based on the phylogeny for 332 yeast species from (Shen *et al*. 2018), except for three species in the MDR clade other than *C. auris*, whose phylogenetic relationships are based on (Muñoz *et al*. 2018). The tip colors show the pathogenic status of the species. The highlighted clades are enriched in known human pathogens. In the table, the first column shows the total number of Hil family homologs per species. The number of homologs that pass each of the three tests for determining their adhesin status are shown in the next three columns. FRV = FungalRV, SP = Signal Peptide and GPI = GPI-anchor. See Materials and Methods for details. The number of homologs passing all three tests is shown in the “final” columns. (B) Boxplots comparing the number of Hil homologs (upper) or the number of putative adhesins passing all three tests (lower) per species between known pathogens and low pathogenic-potential species. Individual species numbers are shown as dots on top of the boxplot. Homologs from *M. bicuspidata* were excluded (see text). Both comparisons are significant at a 0.005 level by either a t-test with unequal variance or Mann-Whitney U test. (C) Maximum likelihood tree based on the Hyphal_reg_CWP domain of the Hil family was constructed using RAxML-NG and corrected with GeneRax based on the species tree. The tree is shown as a cladogram. All 29 homologs in *M. bicuspidata* formed a single group, which is shown as a triangle in dark plum. Homologs from the species in the three highlighted clades in (A) are colored accordingly. CaLo = Candida/Lodderomyces. Homologs from *C. albicans, C. auris* and *C. glabrata* are labeled as Calb, Caur and Cgla, respectively. (D) Species tree showing the number of inferred duplication events on each branch. The gray colors of the tip and internal nodes represent the identified and inferred number of Hil homologs, respectively. The branch color shows the inferred number of duplication events, with 3 or more duplications also shown as a number next to the branch.

Gene duplication and the subsequent functional divergence is a major source for the evolution of novel phenotypes (Zhang 2003; Qian and Zhang 2014; Kuang et al. 2016; Eberlein et al. 2017). In a genome comparison of seven pathogenic *Candida* species and nine low-pathogenic potential relatives, three of the top six pathogen-enriched gene families encode GlycosylPhosphatidylInositol (GPI)-anchored cell wall proteins, namely Hyr/Iff-like, Als-like and Pga30-like (Butler et al. 2009). The first two encode known fungal adhesins (Bailey et al. 1996; Hoyer 2001; Luo et al. 2010). These glycosylated cell wall proteins play key roles in fungal attachment to host endo- and epithelial cells, mediate biofilm formation and iron acquisition, and are well-established virulence factors (HOYER et al. 2008; de Groot et al. 2013; Lipke 2018). It has been suggested that expansion of the cell wall protein repertoire, particularly adhesins, is a key step towards the evolution of yeast pathogens (Gabaldón et al. 2016). This is supported by a study showing that several adhesin families independently expanded in *C. glabrata* and close relatives (Gabaldón et al. 2013). Interestingly, studies of pathogenic *Escherichia coli* found that multiple strains independently acquired genes mediating intestinal adhesion, giving precedence to the hypothesis from a different kingdom (Reid et al. 2000).

Despite the importance of adhesins in both the evolution and virulence of *Candida* pathogens, there is a lack of detailed phylogenetic study for their evolutionary history(Hoyer 2001; Linder and Gustafsson 2008; Gabaldón et al. 2013). Even less is known about their sequence divergence and the role of natural selection in their evolution (Xie et al. 2011). In the newly emerged *Candida auris*, individual adhesins have been characterized but there is little information about their evolutionary relationship with homologs in other *Candida* species and how their sequence diverged (Kean et al. 2018; Singh et al. 2019; Muñoz et al. 2021). In this study we characterized the detail evolutionary history of a yeast adhesin family and used *C. auris* as a focal group to determine how adhesin sequences diverged under various natural selection forces. To choose a candidate adhesin family in *C. auris*, we compared it with the well-studied *C. albicans*, which belongs to the same CUG-Ser1 clade. Of the known adhesins in *C. albicans, C. auris* lacks the Hwp family and has only three Als or Als-like proteins compared with eight Als proteins in *C. albicans* (Muñoz et al. 2018). By contrast, *C. auris* has eight genes with a Hyphal_reg_CWP (PF11765) domain found in the Hyr/Iff family in *C. albicans* (Muñoz et al. 2021). This family was one of the most highly enriched in pathogenic *Candida* species relative to the non-pathogenic ones (Butler et al. 2009). Transcriptomic studies identified two *C. auris* Hyr/Iff-like (Hil) genes as being upregulated during biofilm formation and under antifungal treatment (Kean et al. 2018). Interestingly, isolates from the less virulent *C. auris* Clade II lack five of the eight Hil genes (Muñoz et al. 2021). It is currently not known whether the *C. auris HIL* genes encode adhesins, how they relate to the *C. albicans* Hyr/Iff family genes and how their sequences diverged after duplication.

We show that the Hil family independently expanded multiple times, including in *C. auris* and *C. albicans*. Sequence features and predicted structures of the effector domain indicated that most of the yeast Hil family encode adhesins, including all eight members in *C. auris*, Rates of nonsynonymous-to-synonymous substitutions revealed varying strengths of selective constraint and positive selection acting on the effector domain during the family expansion. Rapid divergence in the repeat-rich central domain led to large variations in length and β-aggregation potential between paralogs and among strains of *C. auris*, likely contributing to phenotypic diversity in adhesion and possibly in virulence.

## Results

### Phylogenetic distribution of the Hyr/Iff-like (Hil) family and its potential to encode adhesin

The Hyr/Iff family was first identified and characterized in *Candida albicans* (Bailey et al. 1996; Richard and Plaine 2007). The family is defined by its N-terminal Hyphally regulated Cell Wall Protein domain (Hyphal_reg_CWP, PF11765), followed by a highly variable central domain rich in tandem repeats (Boisramé et al. 2011). Because the effector domain is more conserved than the repeat region and plays a prominent role in mediating adhesion in known yeast adhesins (Willaert 2018), here we define the Hyr/Iff-like (Hil) family as the group of evolutionarily related proteins sharing the Hyphal_reg_CWP domain, different from a previous definition based on sequence similarity in either the Hyphal_reg_CWP domain or the repeat region (Butler et al. 2009).

To determine the phylogenetic distribution of the Hil family and its association with the pathogenic potential of species, we performed BLASTP searches using the Hyphal_reg_CWP domain from three distantly related Hil homologs as queries (from *C. auris, C. albicans* and *C. glabrata*). We scrutinized the database hits and searched additional assemblies to ensure that their sequences are complete and accurate given the available genome assemblies (Text S1). Using the criteria of *E*-value<10^−5^ and query coverage>50%, we identified a total of 215 proteins containing the Hyphal_reg_CWP domain from 32 species (Fig. 1A, Table S1). No credible hits were identified outside the budding yeast subphylum even after a lower *E*-value cutoff of 10^−3^ was tested, suggesting that this family is specific to this group (Materials and Methods). Species with eight or more Hil family genes fell largely within the Multi-Drug Resistant (MDR) and the Candida/Lodderomyces (CaLo) clades, which include *C. auris* and *C. albicans*, respectively. Only three such species were found outside of the two clades: *C. glabrata, M. bicuspidata* and *K. africana. C. glabrata* is a major opportunistic pathogen that is more closely related to *S. cerevisiae* than to most other *Candida* species (Dujon et al. 2004; Butler et al. 2009; Gabaldón et al. 2013). *M. bicuspidata* is part of the CUG-Ser1 clade. While not a pathogen in humans, it is a parasite of freshwater animals (Hall et al. 2010; Jiang et al. 2022). *K. africana* is not closely related to any known yeast pathogen and its ecology is poorly understood (Gordon et al. 2011).

We then asked how many of the Hil family genes in each species are likely to encode yeast adhesins. To get an initial estimate, we combined a Machine Learning tool for predicting fungal adhesins (Chaudhuri et al. 2011) with predictions for the N-terminal signal peptide and C-terminal GPI-anchor sequence, two features shared by the majority of known fungal adhesins (Lipke 2018). Half of all Hil homologs passed all three tests (Fig. 1A). Notably, *M. bicuspidata* has the largest Hil family among all species, but none of its 29 Hil genes passed all tests. We found most of the identified hits in this species were short relative to the rest of the family (Fig. S1), and 10 of the 29 hits were annotated as being incomplete in the RefSeq database. Further analyses with a better assembled genome and functional studies are needed to determine if the Hil family in this species has unique properties and functions.

### Independent expansion of the Hil family in multiple pathogenic *Candida* lineages

Pathogenic yeast species have on average a larger Hil family and also more of its members were predicted to encode adhesins than in low pathogenic-potential species (Fig. 1B, t-test with unequal variance and Mann-Whitney U test both yielded *P* < 0.005, one-sided test). This naïve comparison doesn’t account for phylogenetic relatedness between species and could result in a false positive association (Levy et al. 2017; Bradley et al. 2018). To address this, we performed phylogenetic logistic regression, which uses the known phylogeny to specify the residual correlation structure among species with shared ancestry (Ives and Garland 2010). We tested for associations between the pathogen status with either the total number of Hil homologs or the number of putative adhesins in each species. Both tests were significant (*P* = 0.005 and 0.007, respectively). Together, these results strongly support an enrichment of the Hil family and the putative adhesins therein among the pathogenic yeast species.

Some adhesin families have undergone independent expansions even among closely related species (Gabaldón et al. 2013). This would result in overestimation of the phylogenetic signal in the above analysis. To further characterize the evolutionary history of the Hil family, including among closely related *Candida* lineages, we reconstructed a species tree-aware maximum likelihood phylogeny for the Hil family based on the Hyphal_reg_CWP domain alignment (Fig. 1C, Fig. S2). We found that homologs from the MDR clade and the Candida / Lodderomyces (CaLo) clade separated into two groups, suggesting that the duplications of the Hil family genes in the two clades occurred independently. To better illustrate the history of gene duplications in the Hil family, we reconciled the gene tree with the species tree and mapped the number of duplications onto the species phylogeny (Materials and Methods). The result showed that the Hil family has independently expanded multiple times, not only between clades but also among closely related species within a clade, such as in *C. albicans* and *C. tropicalis* (Fig. 1D).

### Sequence features of the *C. auris* Hil family support their adhesin status

Experiments have demonstrated that Hil family members function as adhesin in *C. albicans* and more recently for one member in *C. glabrata* (Bailey et al. 1996; Boisramé et al. 2011; Reithofer et al. 2021; Rosiana et al. 2021). To further evaluate the adhesin function of Hil family proteins, we focused on *C. auris*, in which Hil family members were implicated in biofilm formation and response to antifungal treatments, but still remain poorly characterized (Kean et al. 2018). We named the eight *C. auris* Hil family proteins Hil1-Hil8 ordered by their length (Table S2). This differs from the literature, which referred to them by their most closely related Hyr/Iff genes in *C. albicans* (Kean et al. 2018; Jenull et al. 2021; Muñoz et al. 2021). The renaming was to avoid the incorrect implication of one-to-one orthology between the two species (Fig. 1C).

To further assess the adhesin potential for the *C. auris* Hil family, we compared their domain architecture and sequence features to those typical of known yeast adhesins, including a signal peptide, an effector domain, a Ser/Thr-rich and highly glycosylated central domain with tandem repeats and β-aggregation prone sequences and a GPI-anchor signal (Fig. 2A) (de Groot et al. 2013; Lipke 2018). All eight *C. auris* Hil proteins followed this domain architecture (Fig. 2B). Hil1-4 were additionally characterized by an array of regularly spaced β-aggregation prone sequences (red ticks below the protein, Fig. 2B). All eight proteins also had elevated Ser/Thr frequencies in their central domain and were predicted to be heavily O-glycosylated (Fig. 2C). Predicted N-glycosylation was rare except in Hil5 and Hil6 (Fig. 2C). The overall Ser/Thr frequencies in the Hil family proteins were significantly elevated compared with the rest of the proteome (Fig. S3). All eight members were predicted to be fungal adhesins by FungalRV, a support vector machine-based classifier that showed high sensitivity and specificity in eight pathogenic fungi based on sequence features (Chaudhuri et al. 2011).

**Figure 2.**
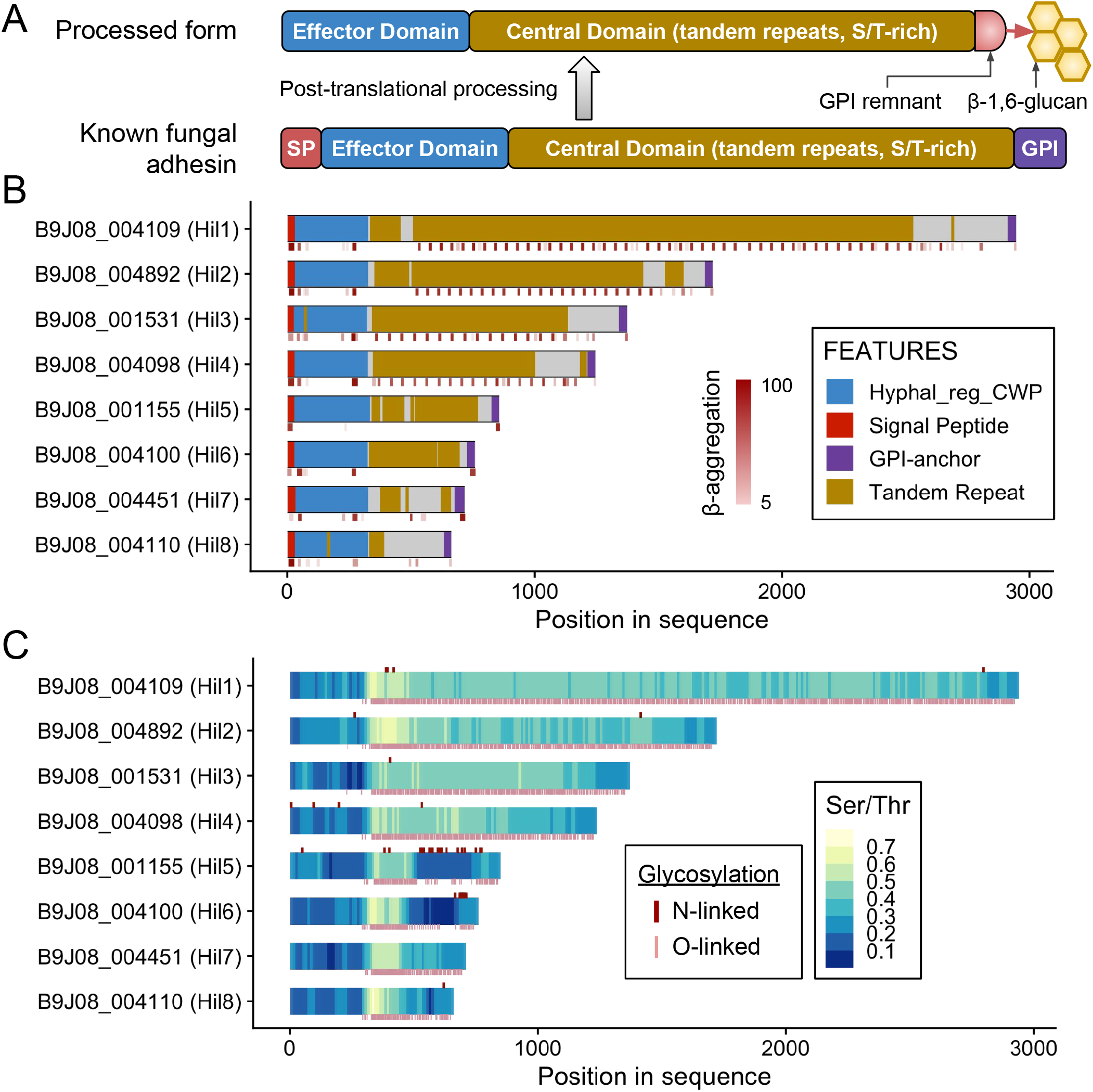
Domain architecture and adhesin-associated features of the *C. auris* Hil family. (A) Diagram depicting the domain organization of a typical yeast adhesin before and after the post-translational processing, adapted from (de Groot *et al*. 2013). (B) Domain features of the eight Hil proteins in *C. auris* (strain B8441). Gene IDs and names are labeled on the left. The short stripes below each diagram are the TANGO predicted β-aggregation prone sequences, with the intensity of the color corresponding to the score of the prediction. (C) Serine and Threonine (Ser/Thr) frequencies in each protein are plotted in 50 aa sliding windows with step size of 10 aa. N-linked and O-linked glycosylation sites were predicted by NetNGlyc 1.0 and NetOGlyc 4.0, respectively, and are shown as short ticks above and below each protein schematic.

### Hyphal_reg_CWP domain in the Hil family is predicted to adopt a β-helical fold similar to unrelated bacterial adhesin binding domains

Crystal structures of the effector domain in several yeast adhesin families, including Als, Epa and Flo, revealed carbohydrate or peptide binding activities supporting the proteins’ adhesin functions (Willaert 2018). The structure of the Hyphal_reg_CWP domain in the Hil family in this study has not yet been experimentally determined. However, crystal structures for the effector domains of two Adhesin-like Wall Proteins (Awp1 and Awp3b) in *C. glabrata*, which are distantly related to those in the Hil family, were recently reported, and the predicted structure of one of *C. glabrata*’s Hil family members (Awp2) was found to be highly similar to the two solved structures (Reithofer et al. 2021). We used AlphaFold2 (Jumper et al. 2021) to predict the structures of the effector domain for two *C. auris* Hil proteins, Hil1 and Hil7 (Fig. 3A, B). Both resemble the *C. glabrata* Awp1 effector domain (Fig. 3C), consisting of a right-handed β-helix at the N-terminus followed by an α-crystallin fold. There are three β-strands in each of the 9 rungs in the β-helix, stacked into three parallel β-sheets (Fig. 3D). The α-crystallin domain consists of seven β-strands forming two antiparallel β-sheets, adopting an immunoglobulin-like β-sandwich fold (Fig. 3E) (Koteiche and Mchaourab 1999; Stamler et al. 2005).

**Figure 3.**
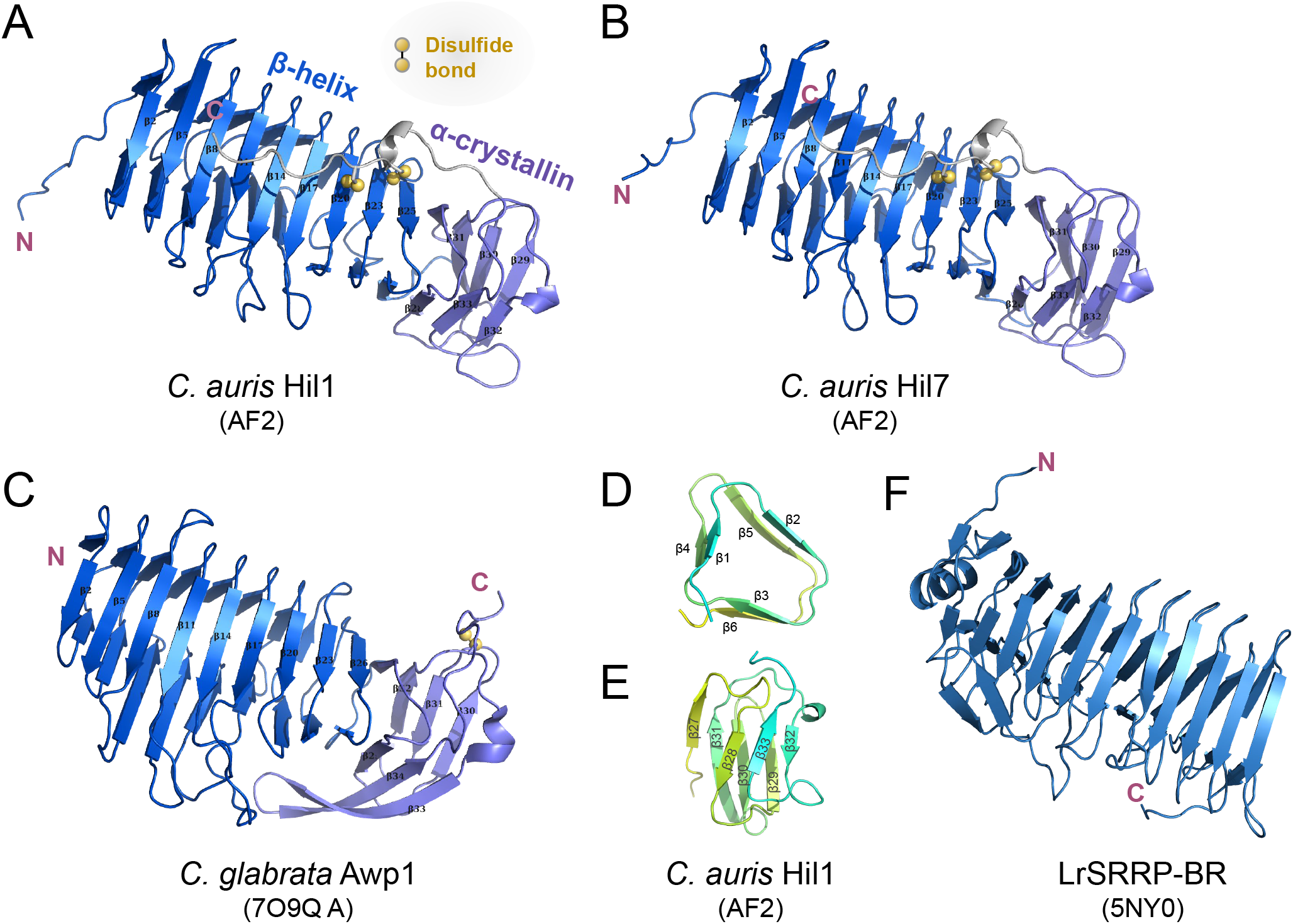
Predicted structures of the Hyphal_reg_CWP domain in two *C. auris* Hil proteins are similar to yeast and bacterial adhesins. (A) and (B) AlphaFold2 (AF2) predicted structures of the Hyphal_reg_CWP domains from *C. auris* Hil1 and Hil7, which consist of a β-helix followed by a α-crystallin domain, with the C-terminal loop linked to the β-helix via two disulfide bonds. (C) Crystal structure of the *C. glabrata* Awp1 effector domain, which is highly similar to *C. auris* Hil1 and Hil7, but with the disulfide bond in a different location. (D) cross-section of the first two rungs of the β-helix in (A), showing the three β-strands per rung. The cyan-to-yellow gradient follows the N- to C-terminus. α-crystallin domain in (A), showing the seven β-strands forming two antiparallel β-sheets. Color is the same as in (D). (F) crystal structure of the Serine Rich Repeat Protein Binding Region (SRRP-BR) from the gram-positive bacterium *L. reuteri*, which adopts a β-helix fold.

The β-strand-rich structure is typical of effector domains in known yeast adhesins, but the β-helix fold at the N-terminus is uncommon (Willaert 2018). Proteins with a β-helix domain often have carbohydrate-binding capabilities and act as enzymes, e.g., hydrolase and pectate lyase (SCOP ID: 3001746). To gain further insight into Hyphal_reg_CWP domain’s function, we searched the PDB50 database for structures similar to what was predicted for *C. auris* Hil1 using DALI (Holm 2022). We identified a number of bacterial adhesins with a highly similar β-helix fold but no α-crystallin domain (Table S3), e.g., Hmw1 from *H. influenzae* (PDB: 2ODL), Tāpirins from *C. hydrothermalis* (PDB: 6N2C), TibA from enterotoxigenic *E. coli* (PDB: 4Q1Q) and SRRP from *L. reuteri* (PDB: 5NY0). For comparison, the binding region of the Serine Rich Repeat Protein 100-23 (SRRP_100-23_) from *L. reuteri* was shown in Fig. 3F (Sequeira et al. 2018). Together, these results strongly suggest that the Hyphal_reg_CWP domain in the *C. auris* Hil family genes mediates adhesion. Additionally, the low sequence identity (12-15%) between the yeast Hyphal_reg_CWP domain and the bacterial adhesins’ binding regions further suggests the two groups have convergently evolved a similar structure to achieve adhesion functions.

### Rapid divergence of the repeat-rich central domain in Hil family proteins in *C. auris*

While the overall domain architecture is well conserved, the eight Hil family proteins in *C. auris* differ significantly in length and sequence of their central domains (Fig. 2B). While not involved in ligand binding, central domains in yeast adhesins are known to play a critical role in mediating adhesion: the length and stiffness of the central domain are essential for elevating and exposing the effector domain (Frieman et al. 2002; Boisramé et al. 2011); and the tandem repeats and β-aggregation sequences within them directly contribute to adhesion by mediating homophilic binding and amyloid formation (Rauceo et al. 2006; Otoo et al. 2008; Frank et al. 2010; Wilkins et al. 2018). Thus, divergence in the central domain of the Hil family has the potential to lead to phenotypic diversity, as shown in *S. cerevisiae* (Verstrepen et al. 2004; Verstrepen et al. 2005).

To determine how the central domain sequences evolved in the *C. auris* Hil family, we used dot plots both to reveal the tandem repeat structure within each protein and to examine the similarity among the paralogs. We found that *C. auris* Hil1 to Hil4 share a ∼44 aa repeat unit, whose copy number varies between 15 and 46, driving their differences in protein length (Fig. 4A). These repeats have conserved periods and sequences (Fig. 4B, Fig. S4). There are two interesting features of this 44 aa repeat unit: a) it contains a heptapeptide “GVVIVTT” that is predicted to be strongly β-aggregation prone, which explains the large number of regularly spaced β-aggregation motifs in Hil1-Hil4 (Fig. 2B); b) it is predicted to form three β-strands in the same orientation (Fig. 4B), raising an interesting question of whether the tandem repeats may adopt a β-structure similar to that of the effector domain. Hil7 and Hil8 encode the same repeat unit but have only one copy (Fig. 4A, red boxes). By contrast, Hil5 and Hil6 encode very different low complexity repeats with a period of ∼5 aa and between 15 to 49 copies (Fig. 4C, D) with relatively low Ser/Thr frequencies (Fig. 2C). Another consequence of encoding only one or zero copy of the 44 aa repeat unit found in Hil1-Hil4 is that Hil5-Hil8 are predicted to have 2-4 β-aggregation prone sequences in contrast to 21-50 in Hil1-Hil4. For comparison, characterized yeast adhesins contain 1-3 such sequences at a cutoff of >30% β-aggregation potential predicted by TANGO (Fernandez-Escamilla et al. 2004; Ramsook et al. 2010; Lipke 2018). The variable lengths, Ser/Thr frequencies and distribution of β-aggregation sequences suggest that the 8 different Hil proteins in *C. auris* are non-redundant, likely playing distinct roles in cell adhesion and other cell-wall related phenotypes.

**Figure 4.**
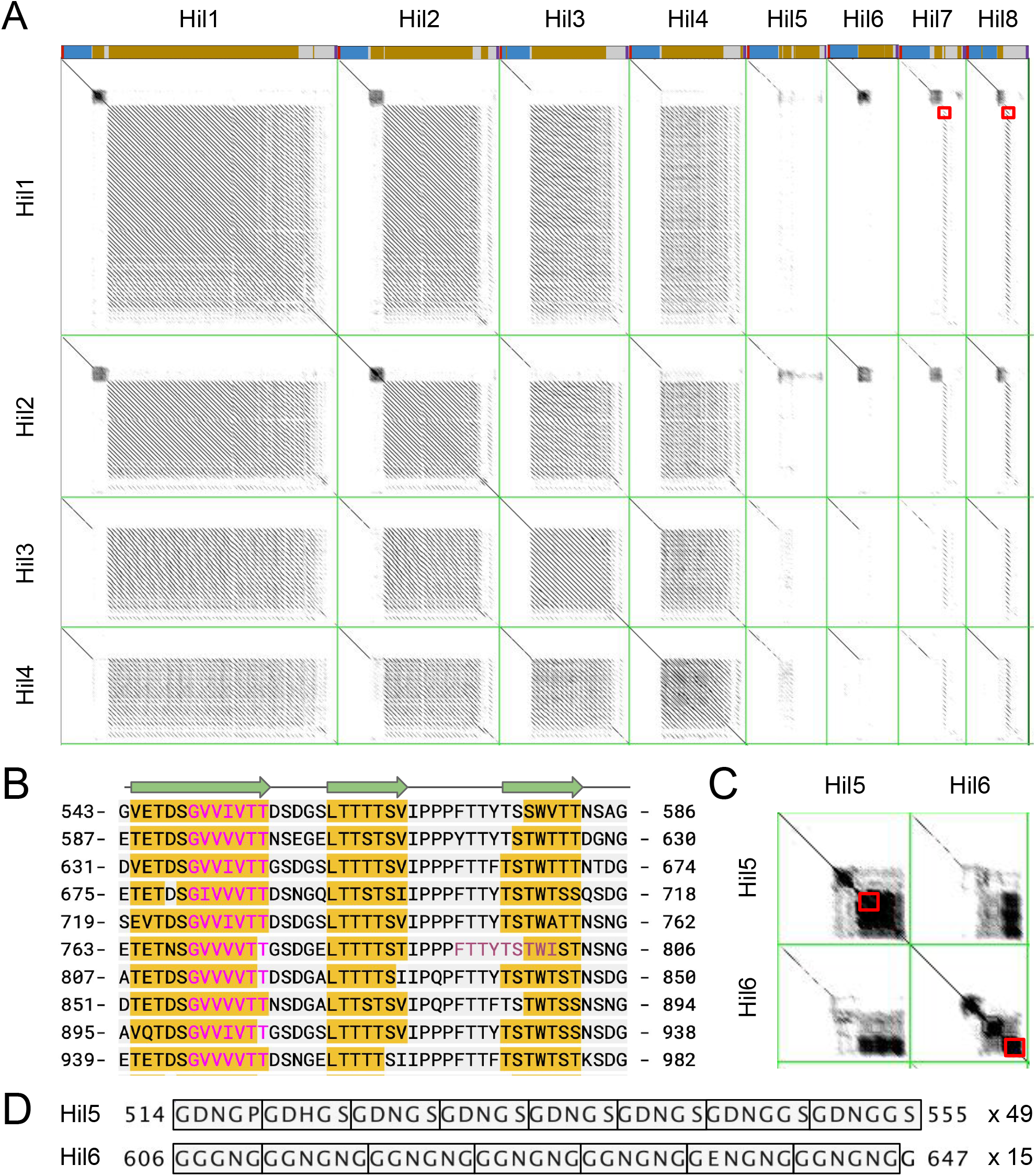
Dotplot shows the tandem repeat structure within and similarity between *C. auris* Hil proteins. (A) Hil1-Hil4 are compared to all eight Hil proteins in *C. auris* including themselves in dotplots (JDotter, Brodie et al 2004) with a sliding window of 50 aa and Grey Map set to 60-245 (min-max). A schematic for each protein is shown above each column (colors same as in Fig. 2). The regions highlighted by the red boxes in row 1 reveal the presence of a single copy of the 44 aa repeat unit in Hil7 and Hil8. (B) Wrapped sequence of aa 543-982 from Hil1 showing the conserved period and sequence of the 44 aa tandem repeat. The magenta and plum fonts indicate motifs predicted by TANGO to have strong (probability > 90%) or moderate (30-90%) β-aggregation potential. The yellow highlighted regions are predicted to form β-strands by PSIPred, with cartoons shown above. (C) Dotplots between Hil5 and Hil6 with the same settings as in (A), showing the low complexity repeats unique to these two. Regions within the two red boxes are shown in (D), with residue numbers shown on both ends. The rectangles delineate individual repeats, with the number of copies for each repeat shown to the right.

Because tandem repeats are prone to recombination-mediated expansions and contractions, we asked if there are variable numbers of tandem repeats (VNTR) among strains in *C. auris*, which could generate diversity in cell adhesive properties as shown in *S. cerevisiae* (Verstrepen et al. 2005). To answer this question, we identified homologs of Hil1-Hil4 in nine *C. auris* strains from three geographically-stratified clades (Muñoz et al. 2018; Muñoz et al. 2021). The genomes of these strains were *de novo* assembled using long-read technologies (Table S4), which allowed us to confidently assess copy number variations within tandem repeats. We identified a total of eight indel polymorphisms in Hil1-Hil4 (Table S5, example alignments in Fig. S5). Except for one 16 aa deletion that is in a single Clade III strain, all seven other indels span one or multiples of the repeat unit and affect all strains within a clade. This is consistent with them being driven by recombination between repeats. The agreement within clades additionally show the indels are not due to sequencing / assembly artifacts, which are not expected to follow the clade labels. As previously reported, Clade II strains lack five of the eight Hil family proteins, including Hil1-4 (Muñoz et al. 2021). Our phylogenetic analysis further showed that this was due to gene losses within Clade II (Fig. S6). The potential relationship between the Hil family size and the virulence profiles of Clade II strains is discussed later.

### Natural selection on the effector domain during the Hil family expansion in *C. auris*

Gene duplication provides raw materials for natural selection and is often followed by a period of relaxed functional constraints on one or both copies, allowing for sub- or neo-functionalization (Zhang 2003; Innan and Kondrashov 2010). Positive selection can be involved in the process, which can lead to a ratio of nonsynonymous to synonymous substitution rates dN/dS > 1 (Yang 1998). Here we ask if the Hyphal_reg_CWP domain in *C. auris* Hil1-Hil8 experienced relaxed selective constraints and/or positive selection following gene duplications, the latter of which would suggest functional diversification. We chose to focus on the Hyphal_reg_CWP domain because of its functional importance and because the high-quality alignment in this domain allowed us to make confident evolutionary inferences (Fig. S7).

Because gene conversion between paralogs can cause distinct genealogical histories for different parts of the alignment and mislead evolutionary inferences (Casola and Hahn 2009), we first identified putatively non-recombining partitions using GARD (Kosakovsky Pond et al. 2006) (Fig. S8), and chose two partitions, P1-414 and P697-981, for maximum-likelihood based analyses using PAML (Yang 2007) (Fig. 5A).

**Figure 5.**
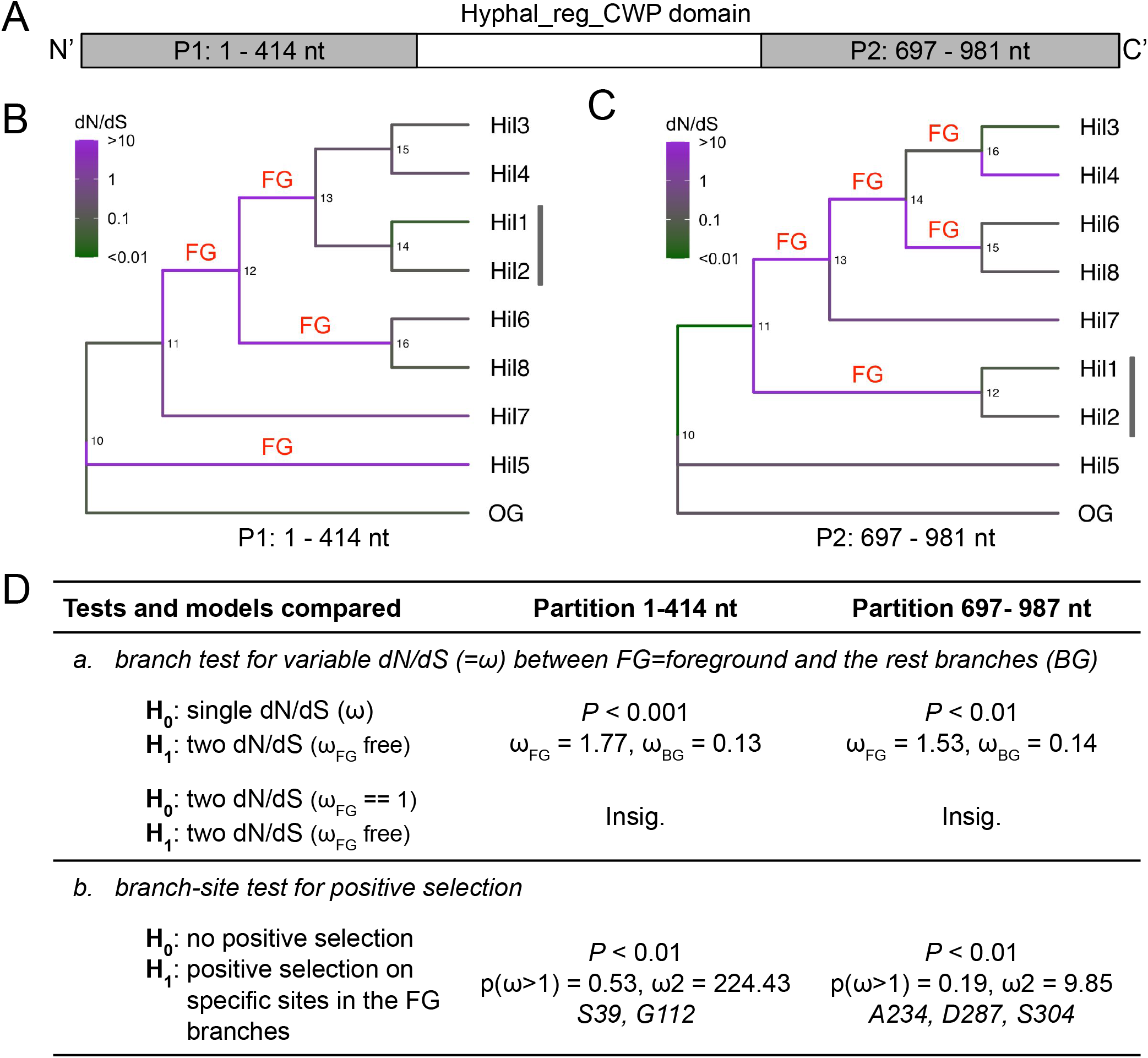
Maximum-likelihood-based analyses for selective pressure variation and role of positive selection on the Hyphal_reg_CWP domain in *C. auris*. (A) Schematic showing the putative non-recombining partitions within the Hyphal_reg_CWP domain determined by GARD (see Fig. S8). The two partitions labeled in gray were studied separately. The numbers refer to the columns in the coding sequence alignment. (B, C) Phylogenetic trees were reconstructed for the two partitions and are shown as a cladogram. The vertical bar next to the Hil1/Hil2 pair indicates the difference in topology between the two trees. Branch colors are based on the dN/dS values estimated from a free-ratio model in PAML. “FG” designate foreground branches, whose dN/dS were greater than 10, except for branch 14..16 in (C), which was selected instead of 16..Hil4 because this would require fewer evolutionary changes in selective forces. We also analyzed the scenario with 16..Hil4 as the foreground and the conclusions remained the same with slightly different *P*-values. (D) Summary of the maximum-likelihood-based tests for selective force heterogeneity and for positive selection. “Insig.” means *P*-value > 0.05. In the branch-site test, p(ω>1) is the total proportion of sites with dN/dS > 1 on the FG branches, and ω2 their estimated dN/dS. The listed sites were identified as being under positive selection with a posterior probability greater than 0.99 by the Bayes Empirical Bayes (BEB). The one-letter code and number refer to the amino acid in the OG sequence and the alignment column (Fig S7).

We first tested if a subset of the sites evolved under positive selection consistently on *all* branches. We found moderate evidence supporting the hypothesis for the P697-981 partition, where the M8 vs. M7 and M8 vs. M8a tests were significant at a 0.01 level, but the conservative test M2a vs M1a was not (Table S6). All three tests were insignificant for the P1-414 partition. Next, we tested for elevated dN/dS on selected branches of the tree, sign of relaxed selective constraints or positive selection. We first estimated the dN/dS for each branch using a free-ratio model and designated those with dN/dS greater than 10 as the “foreground” (Fig. 5B, C, “FG”). We found strong evidence for the FG branches to have a higher dN/dS than the remainder of the tree (log-likelihood ratio test *P* < 0.01, Fig. 5D). There is no evidence, however, for the dN/dS across the entire domain on the FG branches to be greater than 1 (Fig. 5D, *a*, row 2). We then tested the more realistic scenario, where a subset of the sites on the FG branches were subject to positive selection. Using the branch-site test 2 as defined in (Zhang et al. 2005), we found evidence for positive selection on a subset of the sites on the FG branches for both partitions (log-likelihood ratio test *P* < 0.01), and identified residues in both as candidate targets of positive selection with a posterior probability greater than 0.99 (Fig. 5D). We conclude that there is strong evidence for relaxed selective constrain on the Hyphal_reg_CWP domain on some branches following gene duplications; there is also evidence for positive selection acting on a subset of the sites on those branches. However, as the free-ratio model estimates were noisy and the empirical bayes method used to identify the residues under selection lacks power (Zhang et al. 2005) and can produce false positives (Nozawa et al. 2009), the specific branches and residues implicated must be interpreted with caution.

### The yeast Hil family has adhesin-like domain architecture with rapidly diverging central domain sequences

We next examined the entire yeast Hil family to reveal the broader patterns of their evolution. We found that the Hil family in general has elevated Ser/Thr content compared with the rest of the proteome (Fig. S9). Moreover, the majority encode tandem repeats in the central domain (Fig. 6A) and contain predicted β-aggregation prone sequences (Fig. 6B). Together, these features further suggest that a large fraction of the yeast Hil family members likely encode fungal adhesins. While these key features typical of yeast adhesins are conserved, the yeast Hil family exhibits extreme variation in protein length, tandem-repeat content as well as in β-aggregation potential (Fig. 6A, B, S10), extending the pattern seen in *C. auris* (Fig. 2). The length of the protein outside of the Hyphal_reg_CWP domain has a mean ± standard deviation of 822.4±785.8 aa and a median of 608.5 aa. This large variation in protein length is almost entirely driven by the tandem repeats (Fig. 6C, linear regression slope = 1.0, r^2^ = 0.83). A subset of the Hil proteins (vertical bar in Fig. 6A, B) stand out in that 1) they are longer than the rest of the family (1745 vs 770 aa, median protein length) and 2) they have an unusually large number of β-aggregation prone motifs (25 vs 6, median number of TANGO hits per protein). The motifs in this group of proteins are regularly spaced as a result of being part of the tandem repeat unit (median absolute deviation, or MAD, of distances between adjacent TANGO hits less than 5 aa, Fig. 6D). The motif “GVVIVTT” and its variants account for 61% of all hits in this subset and are not found in significant number in the rest of the family. Together, these observations combined with previous experimental studies showing a direct impact of adhesin length and β-aggregation potential on their functions (Verstrepen et al. 2005; Lipke et al. 2012) lead us to propose that the rapid divergence of the Hil family following the parallel expansion led to functional diversification in adhesion in pathogenic yeasts and may have contributed to their enhanced virulence.

**Figure 6.**
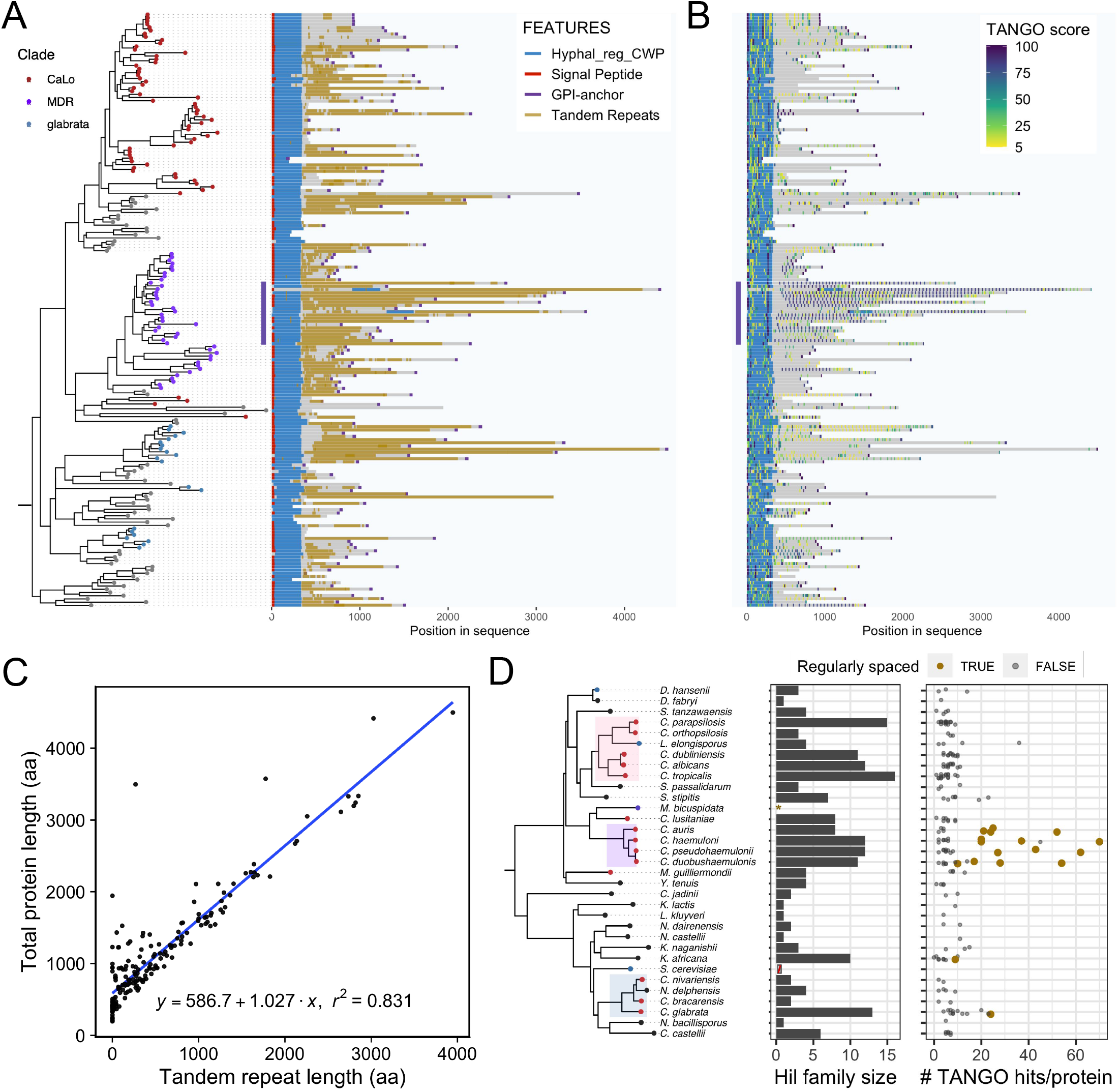
Evolution of protein length and β-aggregation potential in the yeast Hil family. (A) Domain schematic shows that most homologs have a signal peptide at the N-terminus, then the Hyphal_reg_CWP domain and a highly repetitive region central domain, followed by the C-terminal GPI-anchor peptide. Homologs from *M. bicuspidata* were not included because many were annotated as incomplete. They were also excluded from other results in this figure. (B) Distribution of TANGO predicted β-aggregation sequences. The score for each sequence is shown as a color gradient and represents the median of the per-residue probability of aggregation. A vertical bar marks a group of MDR clade sequences that have a large number of β-aggregation prone sequences arranged in regular intervals. (C) X-Y plot showing the relationship between total protein length and tandem repeat sequence length for Hil family homologs. The linear regression line is shown in blue, with coefficients and r^2^ values below. (D) The species tree on the left is the same as in Figure 1. The middle panel shows the number of Hil homologs per species. *M. bicuspidata* homologs were excluded; *S. cerevisiae* was included in the species tree but no Hil homolog was identified in it (see text). The right panel shows the number of predicted β-aggregation prone motifs per Hil homolog. Only motifs with a median probability >= 30% were counted. Proteins are colored in gold if they have five or more such motifs and if the Median Absolute Deviation (MAD) of the inter-motif distances is < 5 aa.

### The yeast Hil family genes are preferentially located near chromosome ends

Several well-characterized yeast adhesin families, including the Flo family in *S. cerevisiae* and the Epa family in *C. glabrata*, are enriched in the subtelomeres (Teunissen and Steensma 1995; De Las Peñas et al. 2003; Xu et al. 2020; Xu et al. 2021). This region is associated with high rates of SNPs, indels and copy number variations, and can undergo ectopic recombination that enables the spread of genes between chromosome ends or their losses (Mefford and Trask 2002; Anderson et al. 2015). To determine if the Hil family is also enriched in the subtelomeric region, we compared their chromosomal locations with the background gene density distribution (Fig. 7A) in species with a chromosomal level assembly (Table S7). To account for the shared evolutionary history, we selected one species per closely related group such that the Hil family homologs in these species were mostly derived through independent duplications based on our gene tree (Fig. S2). The result showed that the Hil family genes are indeed enriched at chromosomal ends (Fig. 7B). A goodness-of-fit test confirmed that the difference between the chromosomal locations of the Hil family and the genome background is highly significant (*P* = 1.3×10^−12^). As ectopic recombination between subtelomeres has been suggested to underlie the spread of gene families (Anderson et al. 2015), we hypothesize that the enrichment of the Hil family towards the chromosome ends is both a cause and consequence of its parallel expansion in different *Candida* lineages.

**Figure 7.**
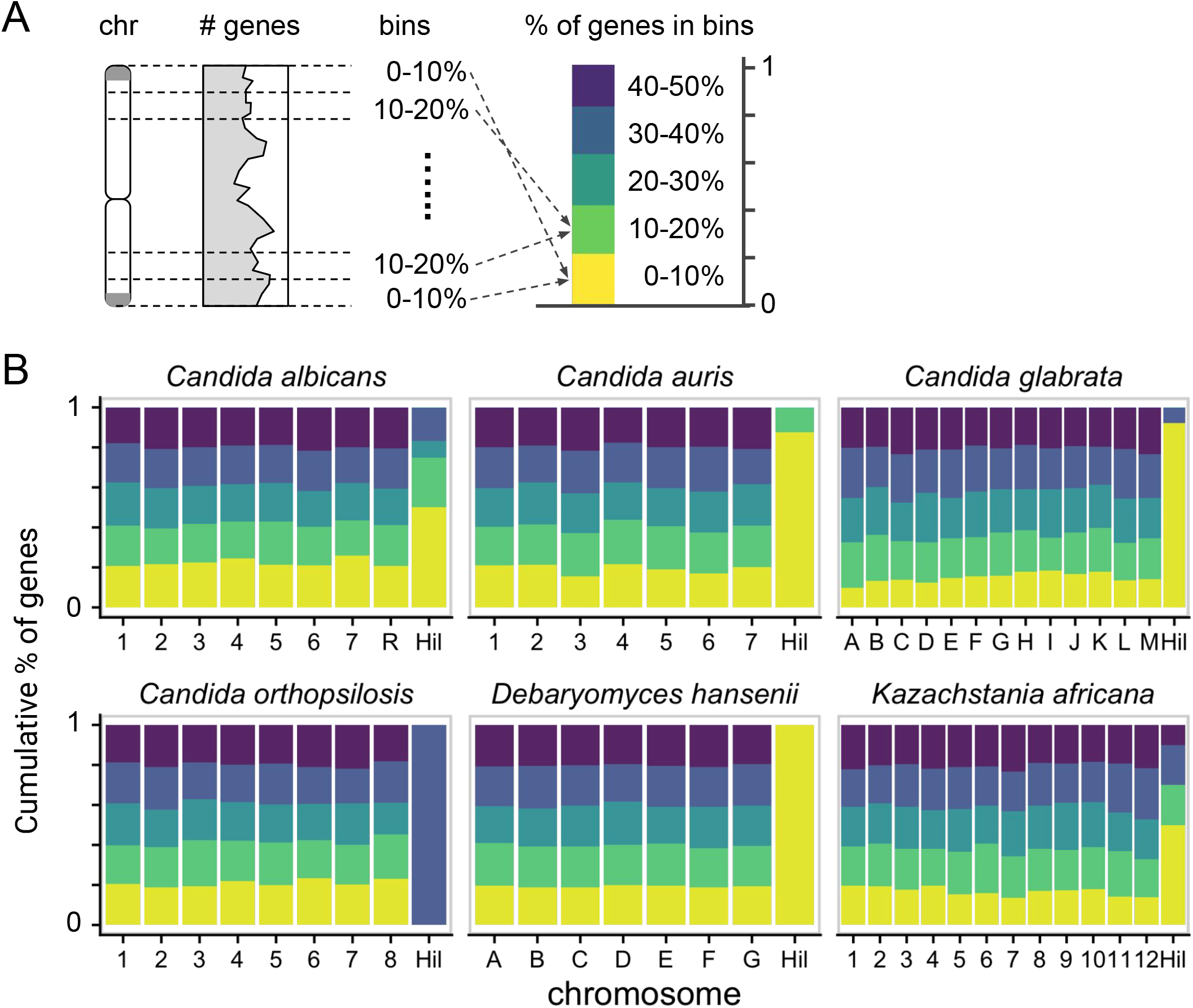
Hil family genes are preferentially located near the chromosome ends. (A) Schematic of the analysis: each chromosome (chr) is folded in half and divided into five equal-length “bins”, ordered by their distance to the nearest telomere. The cumulative bar graph on the right summarizes the gene density distribution in the five bins. (B) Folded gene density distribution for six species with a chromosomal level assembly and more than two Hil family genes. The bin colors are as shown in (A). The Hil homologs in each species are plotted as a separate group. A goodness-of-fit test comparing the distribution of the Hil family genes to the genome background yielded a *P*-value of 1.3×10^−12^.

## Discussion

The repeated emergence of yeast human pathogens in the Saccharomycetes class poses serious health threats, as many emerging pathogenic species are multi-drug resistant or quickly gain resistance (Lamoth et al. 2018; Srivastava et al. 2018). This raises an evolutionary question: are there shared genomic changes in independently derived *Candida* pathogens, which could be key factors behind host adaptation? Yeast adhesin families were among the most enriched gene families in pathogenic lineages relative to the low pathogenic potential relatives (Butler et al. 2009). It has been proposed that expansion of adhesin families could be a key step in the emergence of novel yeast pathogens (Gabaldón et al. 2016). However, detailed phylogenetic studies supporting this hypothesis are rare (Gabaldón et al. 2013), and far less is known about how their sequences diverge and what selective forces are involved during the expansions (Xie et al. 2011; Muñoz et al. 2021). In this study, we found that the Hyr/Iff-like (Hil) family, defined by the conserved Hyphal_reg_CWP domain, is significantly enriched among distantly related pathogenic clades (Fig. 1A, B). This resulted from independent expansion of the family in these clades, including among closely related species (Fig. 1C, D). We also showed that the protein sequences diverged extremely rapidly after duplications, driven mostly by the evolution of the tandem repeats and resulting in large variations in protein length, Ser/Thr content and β-aggregation potential (Fig. 2B, C, Fig. 6). Our evolutionary analyses revealed evidence of relaxed selective constraint and a potential role of positive selection acting on the Hyphal_reg_CWP domain during the family’s expansion in *C. auris* (Fig. 5). We also found the Hil family to be strongly enriched near chromosomal ends (Fig. 7). Overall, our results support the hypothesis that expansion and diversification of adhesin families is a key step towards the emergence of yeast pathogens.

### Genome assembly quality limits gene family evolution studies

Like any study of multi-gene family evolution, our work relies on and is limited by the quality of the genome assemblies. Two additional challenges in our study are due to the fact that Hil family genes are rich in tandem repeats (Fig. 2B, 6A), and many are located near chromosome ends (Fig. 7B), both of which pose problems for genome assemblies. For example, we found significant disagreement in length for 8 of the 16 Hil proteins in *C. tropicalis* between a long-read assembly and the RefSeq assembly, consistent with a recent study (Oh et al. 2020) (Table S8); in *C. glabrata*, we identified 13 Hil family genes in a long-read assembly (GCA_010111755.1) vs 3 in the RefSeq assembly (GCF_000002545.3); 12 of the 13 genes were in the subtelomeres (Xu et al. 2020). However, similar analyses in additional species didn’t reveal these problems, suggesting that the issues were at least in part due to difficulties in some genomes (Text S1). Nonetheless, we acknowledge the possibility of missing homologs and inaccurate sequences, especially in the tandem-repeat region. We thus believe the expected improvements in genome assemblies due to advances in long-read sequencing technologies will be crucial for future studies of the adhesin gene family in yeasts. It is worth noting that our main conclusions about the parallel expansion of the Hil family and its rapid divergence patterns are robust with respect to isolated problems as described above. Also, the long-read technology-based and *de novo* assembled genomes for *C. auris* strains allowed us to confidently assess variation in the Hil family size and tandem repeat copy number between paralogs and among individual strains (Table S4). The accuracy of the tandem-repeat sequences in multiple strains in this species is supported by the conservation of repeat copy numbers within clades (Table S5).

### Evidence for adhesin functions in the Hil family

A few members of the Hil family, e.g., Iff4 in *C. albicans* and Awp2 in *C. glabrata* were shown to mediate adhesiveness to polystyrene (Fu et al. 2008; Kempf et al. 2009; Reithofer et al. 2021). While further experimental studies are needed to establish the adhesin functions of other Hil family members, our work provided bioinformatic support for a large fraction of the Hil family as encoding adhesins (Fig. 2, 6). The predicted β-helix fold of the Hyphal_reg_CWP domain (Fig. 3), while unusual among characterized yeast adhesins (Willaert 2018), is found in many virulence factors residing on the surface of bacteria or viruses as well as enzymes that degrade or modify polysaccharides (Table S3) (Kajava and Steven 2006). The elongated shape and rigid structure of the β-helix are consistent with the functional requirements of adhesins, including the need to protrude from the cell surface and the capacity for multiple binding sites along its length that facilitate adhesion. In a bacterial adhesin – the serine rich repeat protein (SRRP) from the Gram-positive bacterium, *L. reuteri* – a protruding, flexible loop in the β-helix was proposed to serve as a binding pocket for its ligand (Sequeira et al. 2018). Such a feature is not apparent in the predicted structure of the Hyphal_reg_CWP domain. Further studies are needed to elucidate the mechanism of action of this domain and its potential substrates.

The cross-kingdom similarity in adhesin effector domain structure is intriguing in several ways. First, it suggests convergent evolution in bacteria and yeasts. Second, it suggests that what is known about the structure-function relationship in bacteria can provide insight into the Hyphal_reg_CWP domain in yeast. Notably, the *Lr*SRRP shows a pH-dependent substrate specificity that is potentially adapted to distinct host niches (Sequeira et al. 2018). Finally, the similar structure and function of the bacterial and yeast adhesins could mediate cross-kingdom interactions in natural and host environments (Uppuluri et al. 2018).

However, not all Hil family homologs are likely to encode adhesins. Sequence features suggest some Hil family proteins may have non-adhesin functions. For example, 39 of 193 Hil proteins (homologs labeled as incomplete were excluded) have the requisite signal sequence, but lack a GPI anchor attachment site (Fig. S1). One of them, Iff11 in *C. albicans*, was shown to be secreted, and a null mutant of it was found to be hypersensitive to cell wall-damaging agents and less virulent in a murine systemic infection model (Bates et al. 2007). Moreover, 75% of these “SP+, gpi-” proteins are shorter than 600 amino acids, in contrast to only 4% of the 117 proteins having both a signal peptide and a GPI anchor attachment site. Such short, secreted proteins with tandem repeat sequences identical or similar to those present in the cell-wall associated Hil protein counterparts may serve an important regulatory function by bundling with wall associated adhesins as previously suggested for similar subclass of proteins within the Als family (Oh et al. 2019). It is possible that the Hil family has evolved diverse functions broadly related to cell adhesion.

### Ongoing diversification of the Hil family within species

In addition to the parallel expansion and the subsequent rapid sequence divergence in the Hil family between species, we and others also revealed population level variation in both the family size and sequences within *C. auris* (Fig. S5, S6, Table S5) (Muñoz et al. 2021). Notably, among the four geographically stratified clades, Clade II strains lost five of the eight Hil family members (Fig. S6). Besides missing members of the Hil family, Clade II strains also lack seven of the eight members of another GPI-anchor family that is specific to *C. auris* (Muñoz et al. 2021). These coincide with the finding that Clade II strains were mostly associated with ear infections (57/61 isolates according to (Kwon et al. 2019)) rather than hospital outbreaks, as reported for strains from the other clades, and that they were generally less resistant to antifungal drugs (Kwon et al. 2019; Welsh et al. 2019). This raises the question of whether the smaller adhesin repertoire in Clade II strains limits their adhesive capability and results in a different pathology. Similar expansion and contraction of adhesin families have been shown for the *C. glabrata* Hil family (AWP Cluster V) and Epa family (Marcet-Houben et al. 2022), suggesting that dynamic evolution of adhesin families in pathogenic yeasts could be a common pattern. Variations in the tandem repeat copy number in Hil1-Hil4 among *C. auris* strains is also intriguing (Fig. S5). Prior studies of the *S. cerevisiae* Flo proteins have shown that protein lengths directly impact cells’ adhesion phenotypes (Verstrepen et al. 2005) and thus population level variation in adhesin length could further contribute to phenotypic diversity. Lastly, scans for selective sweep in *C. auris* identified Hil and Als family members as being among the top 5% of all genes, suggesting that adhesins are targets of natural selection in the recent evolutionary history of this newly emerged pathogen (Muñoz et al. 2021).

Diversification of adhesin repertoire within a strain can arise from a variety of molecular mechanisms. For example, chimeric proteins generated through recombination between Als family members or between an Als protein’s N terminal effector domain and an Hyr/Iff protein’s repeat region have been shown (Butler et al. 2009; Zhao et al. 2011; Oh et al. 2019). Some of the adhesins with highly diverged central domains may have arisen in this manner (Fig. S10). Gene conversion between members of the same family can also drive the evolution of adhesin families within a species, as shown in *S. cerevisiae* and *C. glabrata* (Verstrepen et al. 2004; Marcet-Houben et al. 2022). Evidence of this in the Hil family was revealed in our analysis of recombination within the effector domain in *C. auris* (Fig. S8).

### Special properties of the central domain in *C. auris* Hil1-Hil4 and related Hil proteins

A subset of Hil proteins represented by *C. auris* Hil1-Hil4 (Fig. 6A, B, vertical bar) stand out in that they are much longer on average and encode a large number of β-aggregation prone sequences compared with the rest of the family (Fig. 6B, D). Behind these properties is a conserved ∼44 aa repeat unit containing a highly β-aggregation prone sequence (“GVVIVTT” and its variants) (Fig. 4B). β-aggregation prone sequences and the amyloid-like interaction they mediate have been extensively studied, especially in the Als protein family in *C. albicans*: they were experimentally shown to mediate aggregation (Otoo et al. 2008; Ramsook et al. 2010) and were crucial for forming protein clusters on cell surfaces known as nanodomains in response to physical tension or sheer forces (Alsteens et al. 2010; Lipke et al. 2012). Recently, they were also shown to mediate cell-cell *trans* interactions via homotypic protein binding (Dehullu et al. 2019; Ho et al. 2019). This may underlie biofilm formation and kin discrimination (Smukalla et al. 2008; Brückner et al. 2020; Lipke et al. 2021). Most known yeast adhesins, including the Als family proteins, encode one to three β-aggregation prone sequences (Ramsook et al. 2010). Unusually, *C. auris* Hil1-Hil4 and their close relatives is that they have as many as 50 such sequences, with each predicted by TANGO to have ∼90% probability of aggregation, whereas the positive threshold for the algorithm is only >5% over 5-6 residues (Fernandez-Escamilla et al. 2004). The structural implications of the vast number of β-aggregation prone motifs may be that such tandem repeat domains are constitutively amyloid in nature, rather than requiring force or other stimuli as in the Als proteins. The functional implications are unclear without the requisite experimental tests. However, we speculate that variations in protein length and β-aggregation potential resulting from the central domain divergence could directly impact the adhesion functions as previously suggested (Verstrepen et al. 2005; Boisramé et al. 2011; Lipke et al. 2012).

### Structural predictions of the tandem repeat region in *C. auris* Hil1 and Hil2

Given the larger number of ∼44 aa repeats in the central domain of *C. auris* Hil1-Hil4 and the prediction that each repeat encodes 3-4 short consecutive β-strands (Fig 4B), we wondered what structural properties this region may have and how it could contribute to the adhesion function. We explored this question using threading based structural prediction tools such as I-TASSER (Yang et al. 2015) and pDOMThreader (Lobley et al. 2009). For the tandem repeat region in the central domain of Hil1, I-TASSER identified (S)-layer protein (SLP) structures (e.g., RsaA from *C. crescentus*, SbsA and SbsC from *G. sterotherophilus*) as among the top structural analogs. These β-strand-rich structures are known to self-assemble to form a 2-dimentional array on the surface of bacteria, mediating a range of functions including adhesion to host cells in pathogens (Fagan and Fairweather 2014). pDOMThreader analyses of the central domains in Hil1 and Hil2 identified a different set of templates, namely bacterial self-associating proteins including Ag43a from uropathogenic *E. coli*, pertactin from *B. pertussis* and the *H. influenzae* hap adhesin. Interestingly, they have β-helical structures like the Hyphal_reg_CWP domain, and the β-helices in them were involved in cell-cell interaction via an interface along the long solenoidal axis for homotypic interactions, which mediate bacterial clumping (Heras et al. 2014) and lead to biofilm formation in *H. influenzae* (Meng et al. 2011). We speculate that the long repeat regions in Hil1 and Hil2 may similarly mediate cell-cell interactions in *C. auris*.

The possibility of the central domains in Hil1 and Hil2 forming a β-helix is interesting in that β-helix is one of the commonly described structural motifs in functional amyloids, e.g., HET-s from the fungus *Podospora anserina* (Wasmer et al. 2008). Such a solenoid-type amyloid is distinguished from other amyloid types in that the β-sheets formed by repeats within the same protein, rather than among distinct monomeric proteins, are suggested to be stabilized not only by polar zippers and hydrophobic contacts, but also by electrostatic interactions between the alternating β-strands (Willbold et al. 2021). Other examples of amyloid forming proteins with a predicted β-helix structure include the imperfect repeat domain in the human premelanosome protein Pmel17 (Louros et al. 2016) and the extracellular curli proteins of Enterobacteriaceae that are involved in biofilm formation and adhesion to host cells (Shewmaker et al. 2009). The proposed solenoidal structure of the central domain of Hil1-Hil4 like proteins, if true, would have two significant implications. First, it confers the necessary rigidity and extended conformation required for cell wall anchored adhesins to extend into the surrounding extracellular milieu. Second, the numerous β-strand rich repeats each containing a highly amyloid prone heptameric sequence, and capable of wrapping into a solenoidal shaped stack, is likely to substantially reduce the rate-limiting nucleation step, which limits the formation of, e.g., an Aβ amyloid fiber. This would allow the formation of extracellular extensions at low protein concentrations without the need for an extensive fiber lengthening process via the incorporation of additional monomeric units. Finally, the observation of solenoid-mediated intercellular interactions in the Hap adhesins suggests that Hil proteins may likewise have a biofilm related function.

### Genomic context

As reported by (Muñoz et al. 2021), we found that the Hil family genes are preferentially located near chromosomal ends in *C. auris* and also in other species examined (Fig. 7). This is similar to previous findings for the Flo and Epa families (Teunissen and Steensma 1995; De Las Peñas et al. 2003; Xu et al. 2020; Xu et al. 2021), as well as the Als genes in some species (Oh et al. 2021). This location bias of the Hil and other adhesin families is likely a key mechanism for their dynamic expansion and sequence evolution via ectopic recombination (Anderson et al. 2015) and by Break-Induced Replication (Bosco and Haber 1998; Sakofsky and Malkova 2017; Xu et al. 2021). Another potential consequence of the Hil family genes being located in subtelomeres is that they may be subject to epigenetic silencing as an additional regulatory mechanism, which can be derepressed in response to stress (Ai et al. 2002). Such epigenetic regulation of the adhesin genes was found to generate cell surface heterogeneity in *S. cerevisiae* (Halme et al. 2004) and lead to hyperadherent phenotypes in *C. glabrata* (Castaño et al. 2005).

### Concluding remarks

To address the lack of candidate adhesins in *C. auris*, we identified and characterized the Hyr/Iff-like (Hil) family in this species and all yeasts. Based on our results, we hypothesize that expansion and diversification of adhesin gene families is a key step towards the evolution of fungal pathogenesis and that variation in the adhesin repertoire contributes to within and between species differences in the adhesive and virulence properties. Future experimental tests of these hypotheses will be important biologically for improving our understanding of the fungal adhesin repertoire, biotechnologically for inspiring additional nanomaterials, and biomedically for advancing the development of *C. auris*-directed therapeutics.

## Materials and Methods

### RESOURCE AVAILABILITY

#### Lead contact

Further information and requests for resources and reagents should be directed to and will be fulfilled by the Lead Contact, Bin Z. He.

#### Data and code availability

All raw data and code for generating the intermediate and final results are available at the GitHub repository at https://github.com/binhe-lab/C037-Cand-auris-adhesin. Upon publication, this repository will be digitally archived with Zenodo and a DOI will be minted and provided to ensure reproducibility.

#### Software and algorithms list

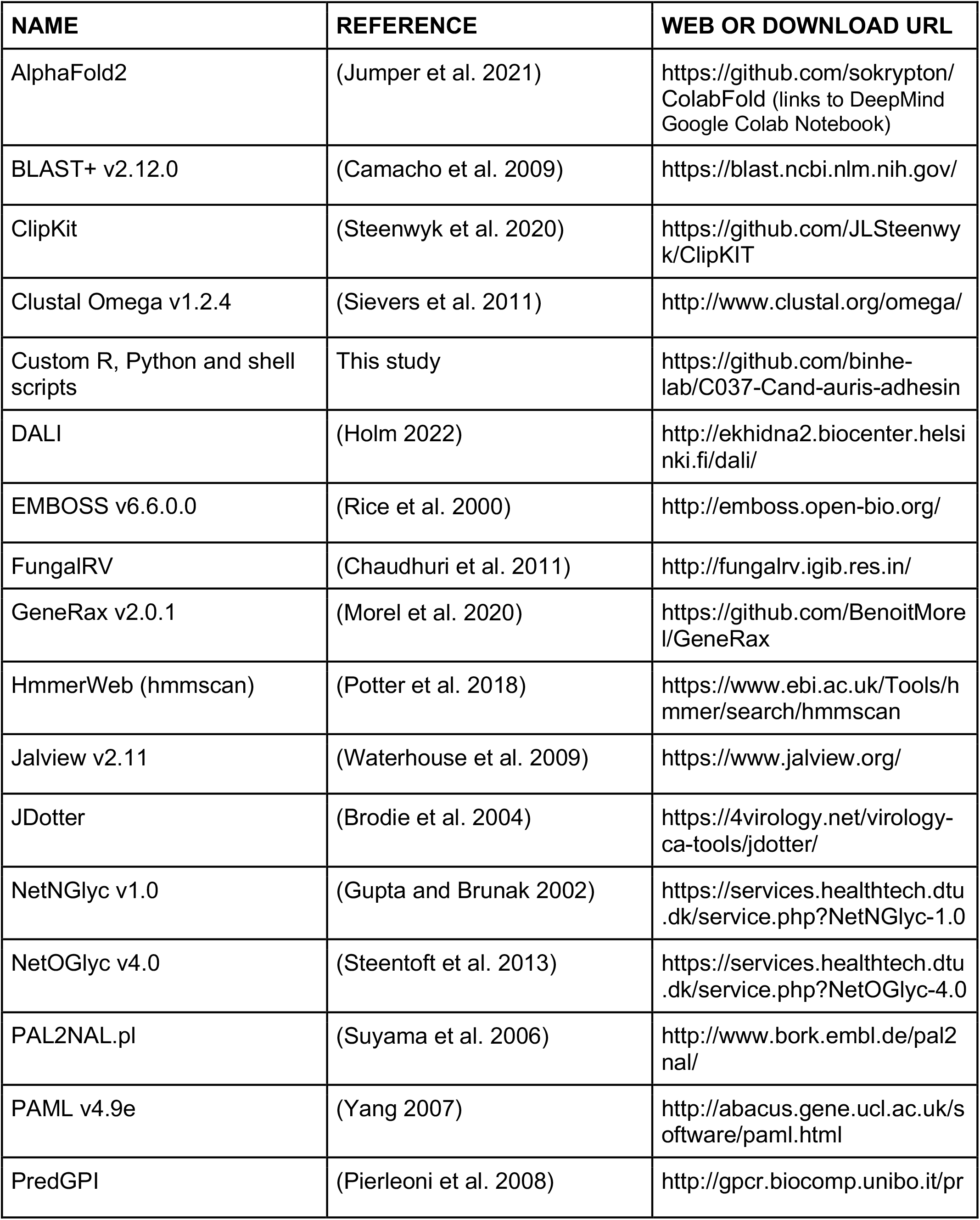
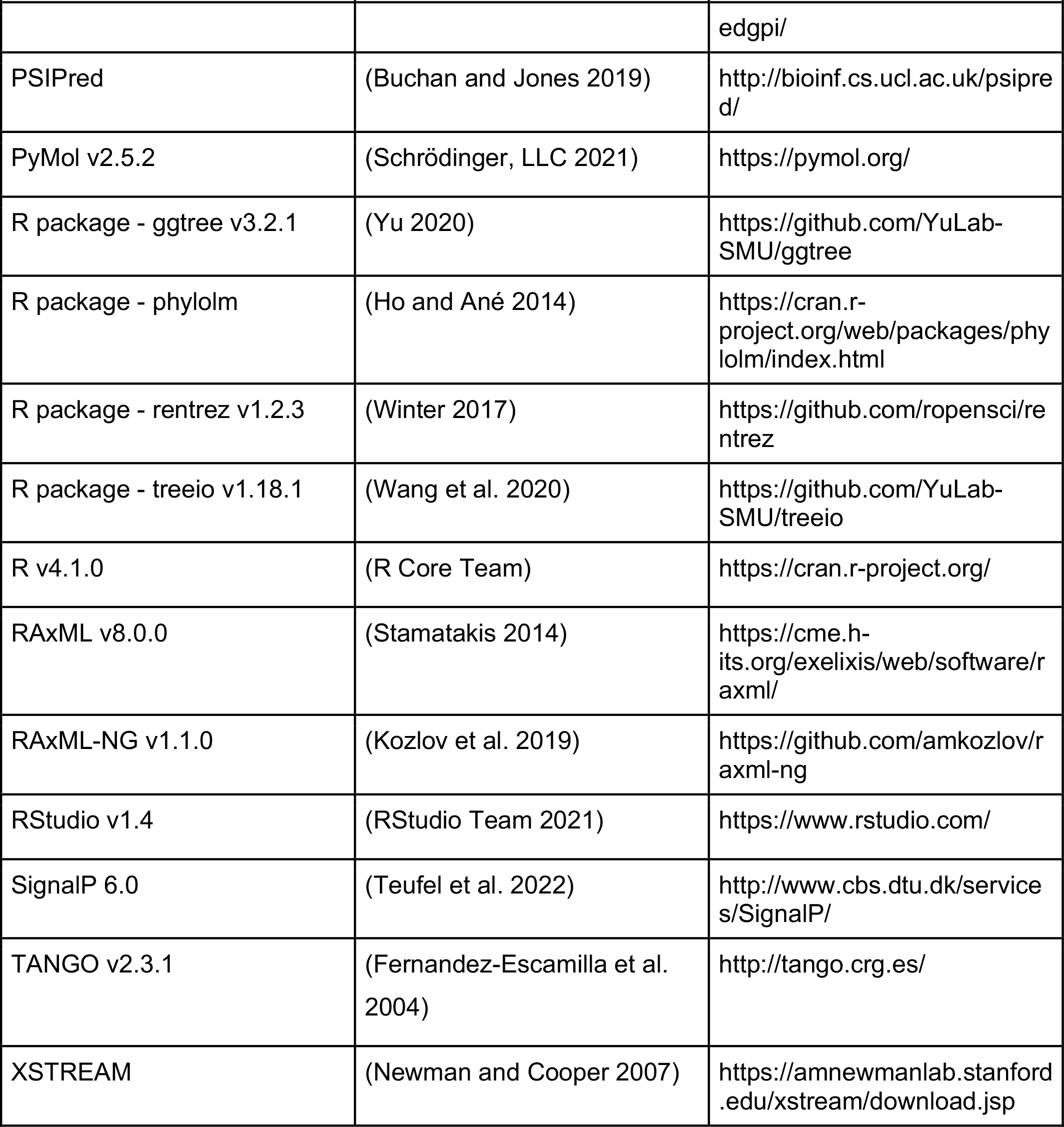

## METHOD DETAILS

### Identify Hyr/Iff-like (Hil) family homologs in yeasts and beyond

To identify the Hil family proteins in yeasts and beyond, we used the Hyphal_reg_CWP domain sequence from three distantly related Hil homologs as queries, namely, *C. albicans* Hyr1 (XP_722183.2), *C. auris* Hil1 (XP_028889033) and *C. glabrata* CAGL0E06600g (XP_722183.2). We performed BLASTP searches in the RefSeq protein database with an *E*-value cutoff of 1×10^−5^, a minimum query coverage of 50% and with the low complexity filter on. All hits were from Ascomycota (yeasts) and all but one were from the Saccharomycetes class (budding yeast). A single hit was found in the fission yeast *Schizosacchromyces cryophilus*. Using that hit as the query, we searched all fission yeasts in the nr protein database, with a relaxed E-value cutoff of 10^−3^ and identified no additional hits. We thus excluded that one hit from downstream analyses. To supplement the RefSeq database, which lacks some yeast species such as those in the Nakaseomyces genus, we searched the Genome Resources for Yeast Chromosomes (GRYC, http://gryc.inra.fr/). Using the same criteria, we recovered 16 additional sequences. To allow for gene tree and species tree reconciliation, we excluded three species that are not part of the 322 species yeast phylogeny (Shen et al. 2018) and not a member of the Multidrug-Resistant clade (Muñoz et al. 2018). Further details, including additional quality control steps taken to ensure that the homolog sequences are accurate and complete, can be found in Text S1. In total, we curated a list of 215 Hil family homologs from 32 species.

### Gene family enrichment analysis

To determine if the Hil family is enriched in the pathogenic yeasts, we performed two analyses. In the first analysis, we divided the species into pathogens vs low-pathogenic potential groups and performed a t-test with unequal variance (also known as Welch’s test) as well as a non-parametric Mann-Whitney U test to compare the Hil family size in the two groups. For both tests, we used either the total size of the family, or the number of putative adhesins as the random variable, and the results were consistent. We excluded homologs from *M. bicuspidata* because 10 of its 29 Hil family proteins were annotated as incomplete in the RefSeq protein database, and also because as a parasite of freshwater crustaceans, it does not fit into either the human pathogen or the low-pathogenic potential group. *S. cerevisiae* was included in the comparison as an example of species with zero members of the Hil family. We chose *S. cerevisiae* because we could be confident about its lack of a Hil family homolog thanks to its well assembled and well annotated genome.

In the second test, we used phylogenetic logistic regression (Ives and Garland 2010) to account for the phylogenetic relatedness between species. We used the `phyloglm` function in the `phylolm` package in R, with {method = “logistic_IG10”, btol = 50, boot = 100}. The species tree, including the topology and branch lengths, were based on the 322 species phylogeny from (Shen et al. 2018), supplemented by the phylogenetic relationship for the MDR clade based on (Muñoz et al. 2018). The *P*-values based on phylogenetically specified residual correlations were reported.

### Phylogenetic analysis of the Hil family and inference of gene duplications and losses

To infer the evolutionary history of the Hil family, we reconstructed a maximum-likelihood tree based on the alignment of the Hyphal_reg_CWP domain. First, we used hmmscan (HmmerWeb version 2.41.2) to identify the location of the Hyphal_reg_CWP domain in each Hil homolog. We used the “envelope boundaries” to define the domain in each sequence, and then aligned their amino acid sequences using Clustal Omega with the parameter {--iter=5}. We then trimmed the alignment using ClipKit with its default smart-gap trimming mode (Steenwyk et al. 2020). RAxML-NG v1.1.0 was run in the MPI mode with the following parameters on the alignment: “raxml-ng-mpi --all --msa INPUT --model LG+G --seed 123 --bs-trees autoMRE”. The resulting tree was corrected using GeneRax, which seeks to maximize the joint likelihood of observing the alignment given the gene family tree (GFT) and observing the GFT given the species phylogeny, using the parameter {--rec-model UndatedDL}. The species tree used is the same as the one used for the phylogenetic logistic regression above. In addition to correcting the gene family tree, GeneRax also reconciled it with the species tree and inferred duplication and loss event counts on each branch. Tree annotation and visualization were done in R using the treeio and ggtree packages (Wang et al. 2020; Yu 2020).

To infer the phylogenetic tree for the Hil family homologs in various *C. auris* strains and infer gains and losses within species, we identified orthologs of the *HIL* genes in representative strains from the four major clades of *C. auris* (B8441, B11220, B11221, B11243) (Muñoz et al. 2018). Orthologs from two MDR species, *C. haemuloni* and *C. pseudohaemulonis*, and from *D. hansenii* were included to help root the tree. The gene tree was constructed as described above. To root the tree, we first inferred a gene tree without the outgroup (*D. hansenii*) sequences in the alignment. Then, the full alignment with the outgroup sequences along with the gene tree from the first step were provided to RAxML to run the Evolutionary Placement Algorithm (EPA) algorithm (Berger et al. 2011), which identified a unique root location. To reconcile the gene tree with the species tree, we performed maximum likelihood-based gene tree correction using GeneRax (v2.0.1) with the parameters: {--rec-model UndatedDL}. The species tree was based on (Muñoz et al. 2018).

### Prediction for fungal adhesins and adhesin-related sequence features

**1)** The potential of Hil homologs encoding fungal adhesins was assessed using FungalRV, a Support Vector Machine-based fungal adhesin predictor (Chaudhuri et al. 2011). Proteins passing the recommended cutoff of 0.511 were considered positive. **2)** Signal Peptide was predicted using the SignalP 6.0 server, with the “organism group” set to Eukarya. The server reported the proteins that had predicted signal peptides. No further filtering was done. **3)** GPI-anchor was predicted using PredGPI using the General Model. Proteins with a false positive rate of 0.01 or less were considered as containing a GPI-anchor. **4)** Tandem repeats were identified using XSTREAM with the following parameters: {-i.7 -I.7 -g3 -e2 -L15 -z -Asub.txt -B - O}, where the “sub.txt” was provided by the software package. **5)** β-aggregation prone sequences were predicted using TANGO v2.3.1 with the following parameters: {ct=“N” nt=“N” ph=“7.5” te=“298” io=“0.1” tf=“0” stab=“-10” conc=“1” seq=“SEQ”}. **6)** Serine and Threonine content in proteins were quantified using `freak` from the EMBOSS suite, with a sliding window of 100 or 50 aa and a step size of 10 aa. To compare with proteome-wide distribution of Ser/Thr frequency, the protein sequences for *C. albicans* (SC5314), *C. glabrata* (CBS138) and *C. auris* (B11221) were downloaded from NCBI Assembly database (IDs in Table S7) and the frequency of serine and threonine residues were counted for each protein. **7)** O-linked and N-linked glycosylations were predicted using NetOGlyc (v4.0) and NetNGlyc (v1.0) servers.

### Structural prediction and visualization for the Hyphal_reg_CWP domain

To perform structural predictions using AlphaFold2, we used the Google Colab notebook (https://colab.research.google.com/github/deepmind/alphafold/blob/main/notebooks/AlphaFold.ipynb) authored by the DeepMind team. This is a reduced version of the full AlphaFold version 2 in that it searches a selected portion of the environmental BFD database, and doesn’t use templates. The Amber relaxation step is included, and no other parameters other than the input sequences are required. DALI was used to search for similar structures in the PDB50 database. Model visualization and annotation were done in PyMol v2.5.2. Secondary structure prediction for *C. auris* Hil1’s central domain was performed using PSIPred.

### Dotplot

To determine the self-similarity and similarity between the eight *C. auris* Hil proteins, we made dotplots using JDotter (Brodie et al. 2004). The window size and contrast settings were labeled in the legends for the respective plots. The self-alignment for *C. auris* Hil1 tandem repeats was visualized using Jalview v2.11.

### Identification of intraspecific tandem repeat copy number variations among *C. auris* strains

To identify polymorphisms in Hil1-Hil4 in diverse *C. auris* strains, we downloaded the genome sequences for the following strains from NCBI: Clade I - B11205, B13916; Clade II - B11220, B12043, B13463; Clade III - B11221, B12037, B12631, B17721; Clade IV - B11245, B12342 (Table S4). The amino acid sequences for Hil1-Hil4 from the strain B8441 were used as the query to search the nucleotide sequences of the above assemblies using TBLASTN, with the following parameters {-db_gencode 12 -evalue 1e-150 -max_hsps 2}. Orthologs in each strain were curated based on the BLAST hits to either the Hyphal_reg_CWP domain alone or the entire protein query. All Clade II strains had no hits for Hil1-Hil4. Several strains in Clade I, III and IV were found to lack one or more Hil proteins (Table S5). But upon further inspection, it was found that they had significant TBLASTN hits for part of the query, e.g., the central domain, and the hits were located at the end of a chromosome, suggesting the possibility of incomplete or misassembled sequences. Further experiments will be needed to determine if those *HIL* genes are present in those strains.

### Estimation of dN/dS ratios and model comparisons

We used ‘codeml’ in PAML (v4.9e) to perform evolutionary inferences on the Hyphal_reg_CWP domain in *C. auris*. We first used Clustal Omega to align the amino acid sequences for the Hyphal_reg_CWP domain from Hil1-Hil8 from *C. auris* similar to how we generated the multiple sequence alignment for all Hil proteins. A closely related outgroup (XP_018709340.1 from *M. bicuspidata*) was included to root the tree. We then generated a coding sequence alignment from the protein alignment using PAL2NAL (Suyama et al. 2006). We used GARD (Kosakovsky Pond et al. 2006) to analyze the coding sequence alignment to detect gene conversion events. The web service of GARD on datamonkey.org was run with the following parameters: {data type: nucleotide, run mode: normal, genetic code: yeast alternative nuclear, site-to-site rate variation: general discrete, rate classes: 3}. Based on the results, we identified two putatively non-recombining partitions, P1 = 1-414 and P2 = 697-981 (the numbers refer to the alignment columns). We then separately analyzed the two partitions in PAML. To test hypotheses about positive selection on a subset of the sites on all branches, we compared models M2a vs M1a, M8 vs M7 and M8a vs M8. The first 4 models were specified by: {seqtype = 1, CodonFreq = 1, model = 0, NSsites = 0,1,2,7,8, icode = 8, fix_kappa = 0, kappa = 2, fix_omega = 0, omega = 0.4, cleandata = 1}. The model M8a is additionally specified by { seqtype = 1, CodonFreq = 1, model = 0, NSsites = 8, fix_omega = 1 and omega = 1, cleandata = 1}. To test hypotheses for variable dN/dS on different branches (no variation across sites), we used {model = 0 or 1 or 2, NSsites = 0}, with the rest being the same as the site tests. Model = 0 specified the single ratio model, model = 1 the free ratio model and model = 2 the user-defined model. For the user-defined model, we first used estimates from the free ratio model to designate a set of branches with dN/dS > 10 as the foreground and then tested if their dN/dS was significantly different from the rest of the tree by comparing a two-ratio model with the single-ratio model. Since the results were significant, we further tested if the foreground dN/dS was significantly greater than 1, by comparing the two-ratio model to a constrained version of the model where omega was fixed at 1. For branch-site test, we used {model = 2, NSsites = 2, fix_omega = 0, omega = .4} as the alternative model and {model = 2, NSsites = 2, fix_omega = 1, omega = 1} as the null to test for positive selection on a subset of the sites on the foreground branches. Sites under positive selection were identified using the Bayes Empirical Bayes (BEB) procedure, with a posterior probability threshold of 0.99.

### Chromosomal locations of Hil family genes

To compare the chromosomal locations of the Hil family genes to the background distribution, we selected eight species whose genomes were assembled to a chromosomal level and are not within a closely related group, including *C. albicans, D. hansenii, C. orthopsilosis, K. africana, K. lactis, N. dairenensis, C. auris* and *C. glabrata* (Table S7). We did not include some species, e.g., *C. dubliniensis*, to minimize statistical dependence due to shared ancestry. The RefSeq assembly for *C. auris* was included even though it was at a scaffold level because a recent study showed that seven of its longest scaffolds were chromosome-length, allowing the mapping of the scaffolds to chromosomes (Muñoz et al. 2021, Supplementary Table 1). To determine the chromosomal locations of the Hil homologs in these eight species, we used Rentrez v1.2.3 (Winter 2017) in R to retrieve their chromosome ID and coordinates. To calculate the background gene density on each chromosome, we downloaded the feature tables for the eight assemblies from the NCBI assembly database and calculated the location of each gene as its start coordinate divided by the chromosome length. To compare the chromosomal location of the Hil family genes to the genome background, we divided each chromosome into five equal-sized bins based on the physical distance to the nearest chromosomal end. We calculated the proportion of genes residing in each bin for the Hil family or for all protein coding genes. To determine if the two distributions differ significantly from one another, we performed a goodness-of-fit test using either a Log Likelihood Ratio (LLR) test or a Chi-Squared test, as implemented in the XNomial package in R (Engels 2015). The LLR test *P*-value was reported.

## Acknowledgement

We thank members of the Gene Regulatory Evolution lab for discussion. We thank Yong Zhang (CAS Zoology Institute), Peter Lipke (Brooklyn College), John McCutcheon (ASU), Matthew Hahn (Indiana University), Rich Lenski (MSU), Josep Comeron (University of Iowa), Dan Weeks (University of Iowa), Lois Hoyer (UIUC) and two other anonymous reviewers for providing useful suggestions and critical comments. We apologize to those who have provided help and whose names were not mentioned. Dr. Bin Z. He is supported by NIH R35GM137831. Lindsey F. Snyder was supported by the NIH Predoctoral Training grant T32GM008629. Rachel A. Smoak is supported by an NSF Graduate Research Fellowship Program under Grant No. 1546595, with additional support through the NSF Division of Graduate Education under Grant No. 1633098.

---

**Supplementary Figure 1.**
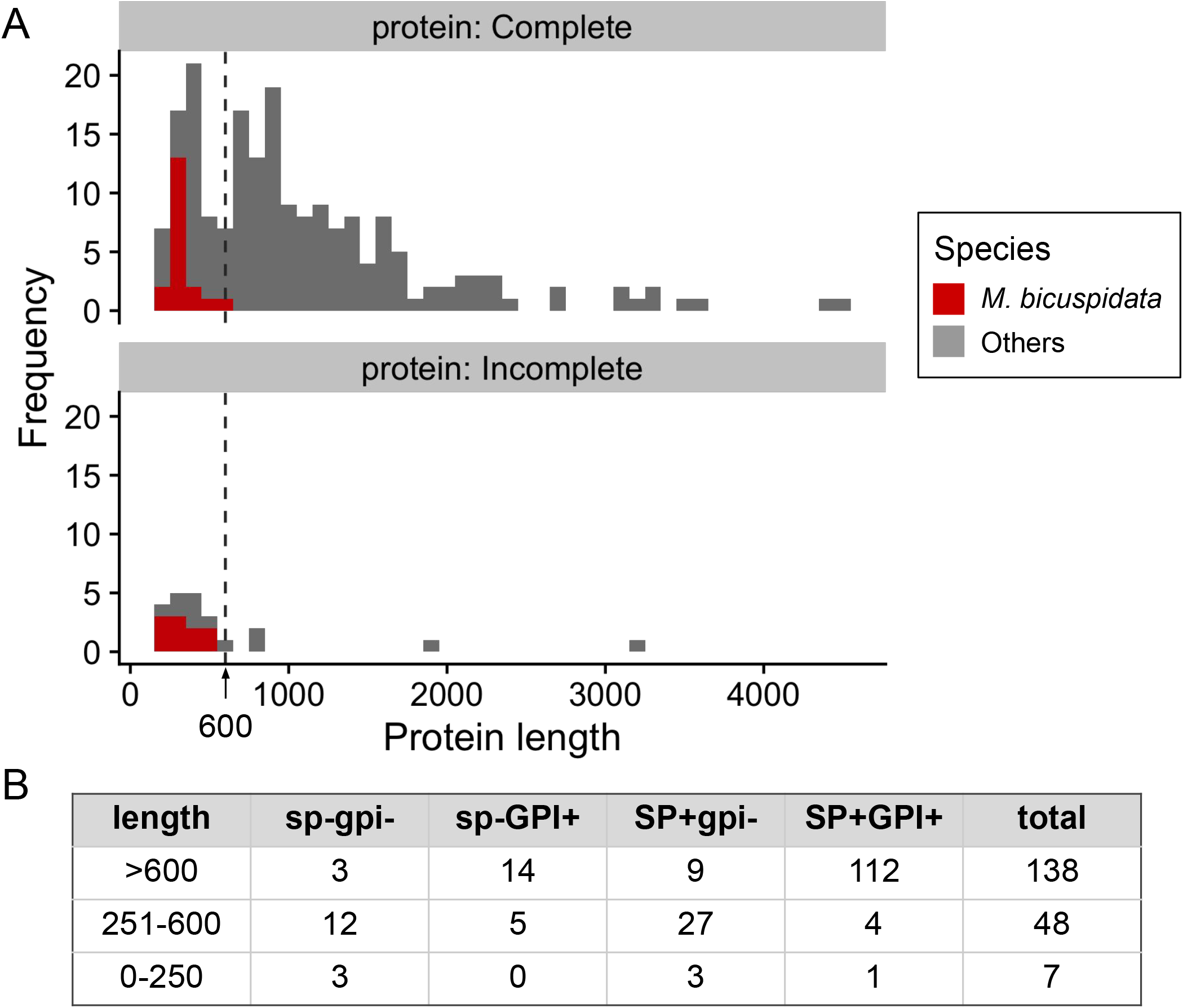
Hil family proteins’ length distribution and grouping by signal peptide (SP) and GPI-anchor signal presence. (A) Histogram showing the distribution of protein lengths for Hil family proteins from 32 yeast species. Top: protein sequence records were labeled as complete or “NA”; bottom: proteins labeled as incomplete (no-right, no-left, no-ends). Most of the short sequences (<600 aa, dashed vertical line) came from the species *M. bicuspidata* (red) (B) Summary of the number of Hil family proteins predicted to have a signal peptide (SP+) and GPI-anchor signal (GPI+), grouped by protein length. Proteins labeled as incomplete were excluded from this table.

**Supplementary Figure 2.**
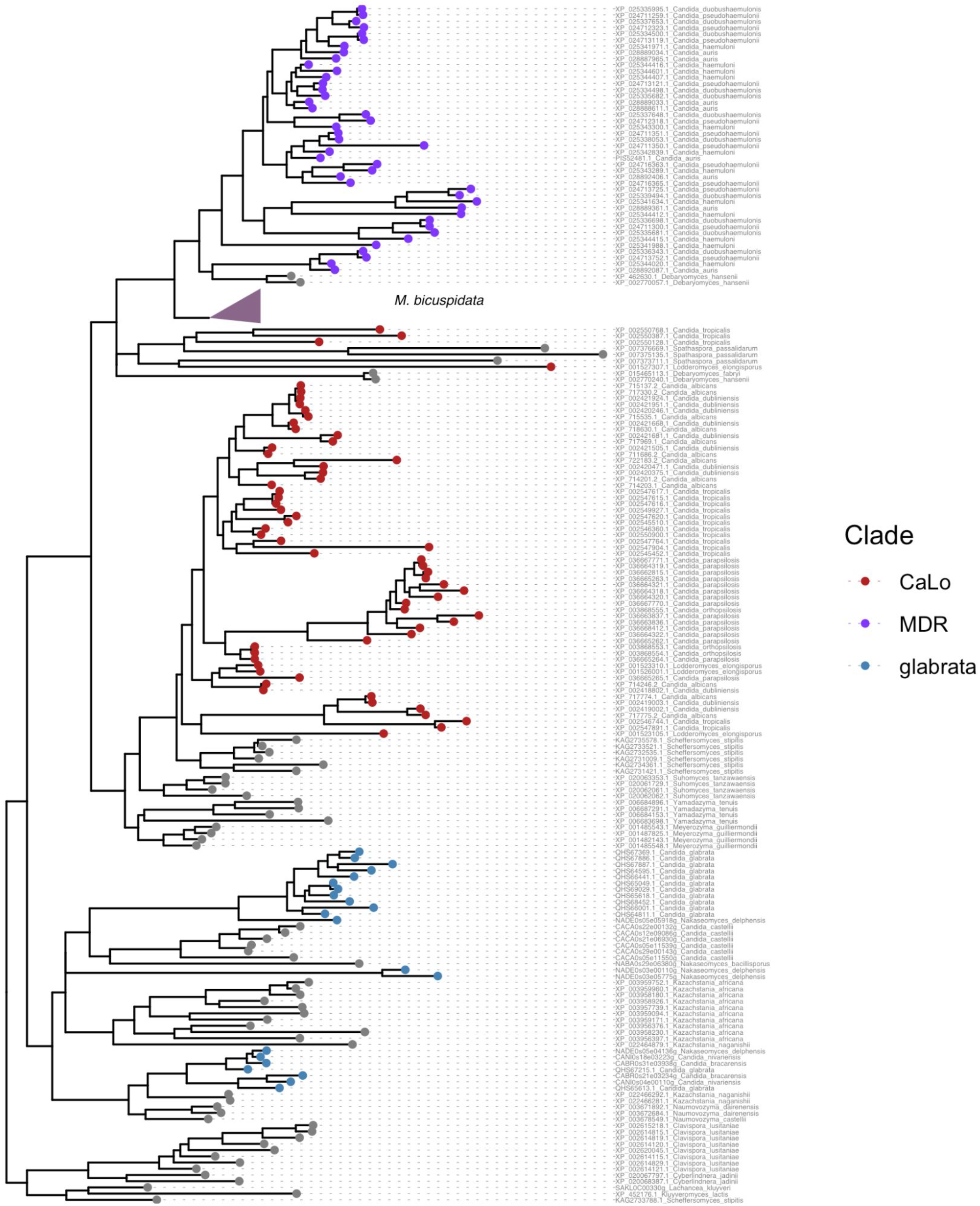
Maximum likelihood tree for the Hil family genes. This tree is identical to the one shown in Fig 1C, except that it is shown in a rectangular format with the sequence names in the form of refseqID_species_name. Note that all 29 *M. bicuspidata* homologs form a single clade, which is collapsed for the ease of viewing. Their sequence IDs can be found in Table S1.

**Supplementary Figure 3.**
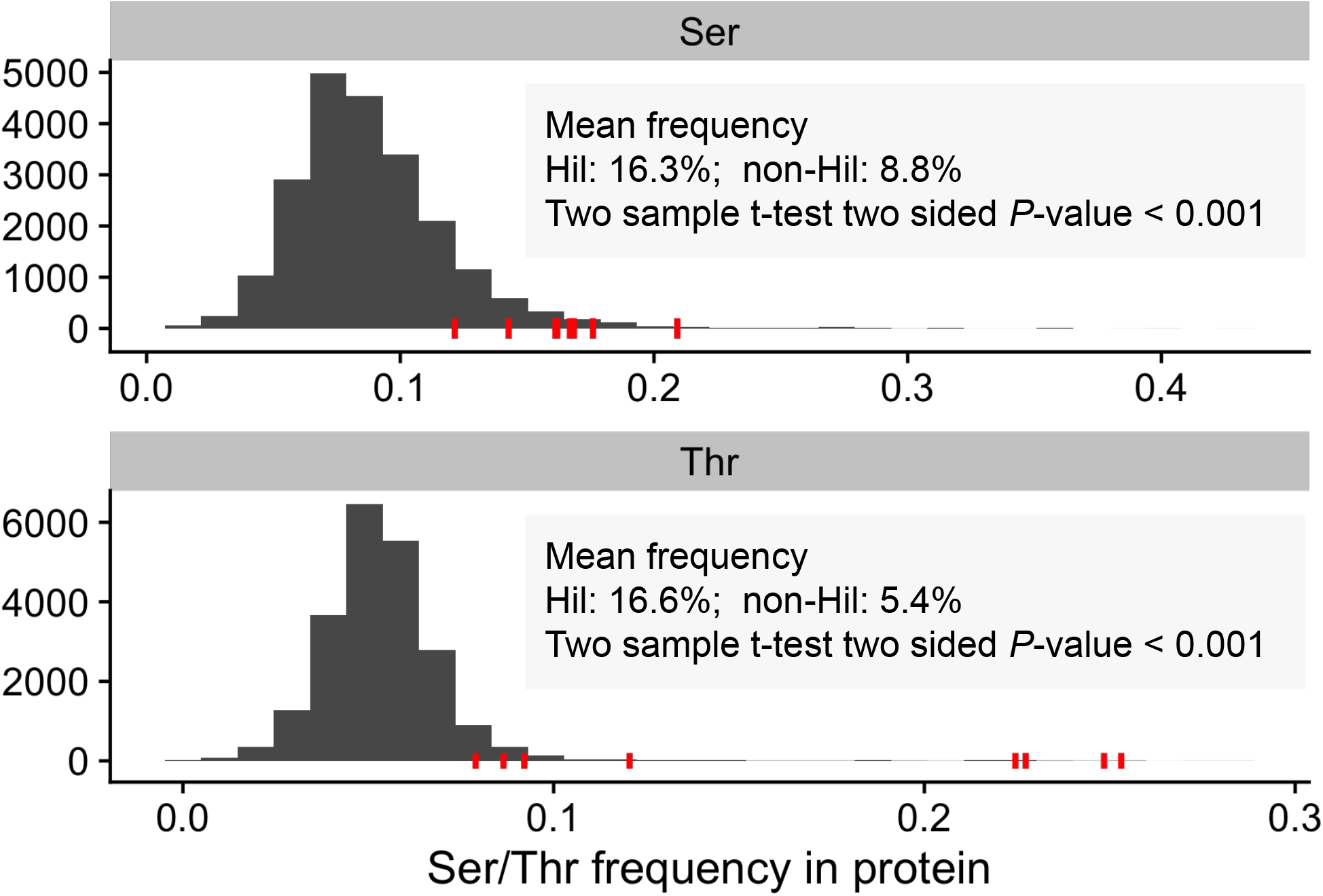
Comparison of the Ser/Thr frequencies in *C. auris* Hil family members with all protein-coding genes in *C. auris*. B8441 strain genome is used for this analysis. The frequency of Ser or Thr residues as a percent of the entire protein length is plotted as a histogram for all protein-coding genes. Red ticks indicate the eight Hil genes. A student’s t-test was used to assess the significance of the difference in Ser/Thr frequencies between the Hil family proteins vs the rest of the proteome.

**Supplementary Figure 4.**
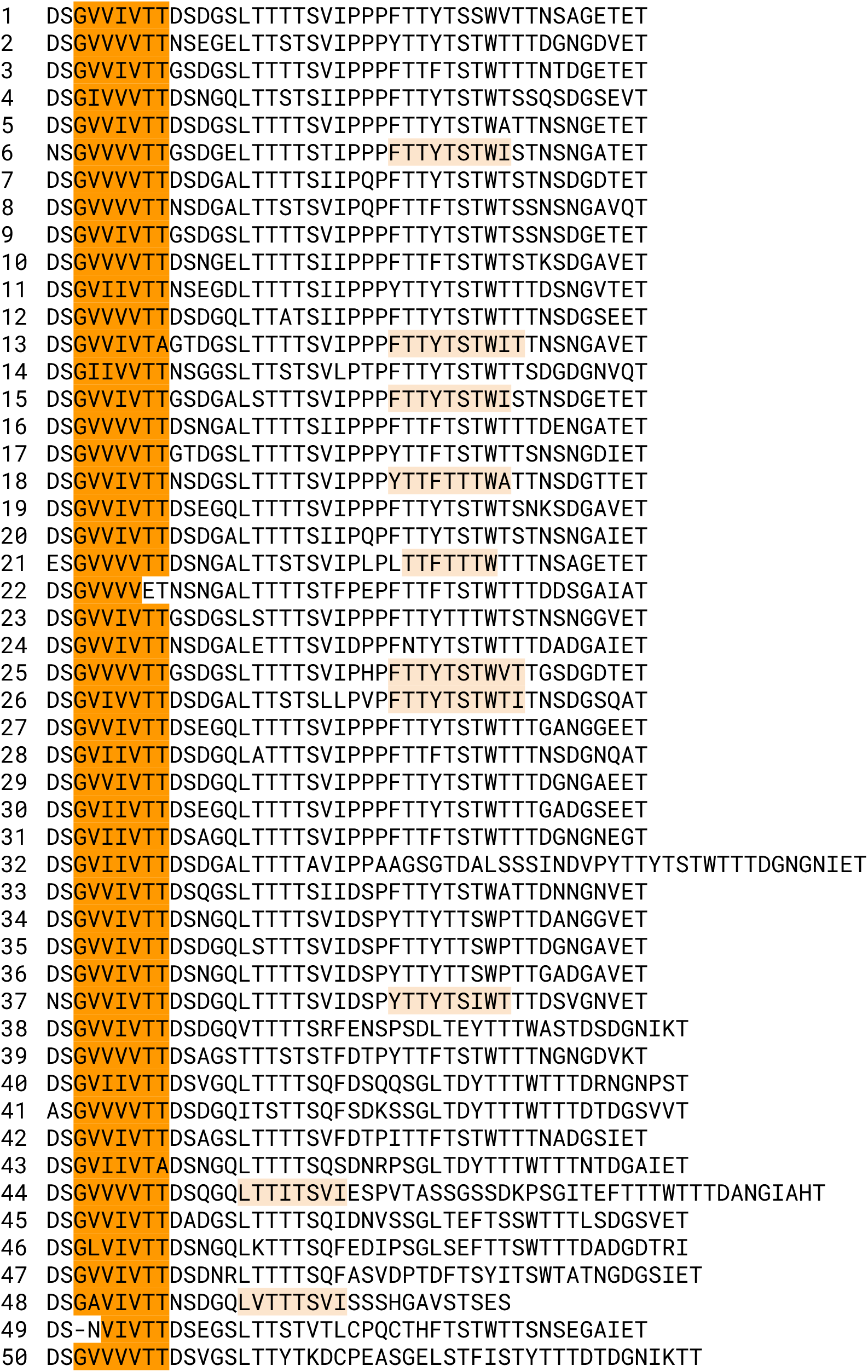
Tandem repeats in the *C. auris* Hil1 central domain. The majority of the 50 tandem repeat copies have a conserved 44 aa period. Dark and light orange highlights show sequences predicted by TANGO to have strong (>90%) or moderate (30-90%) β-aggregation potentials.

**Supplementary Figure 5.**
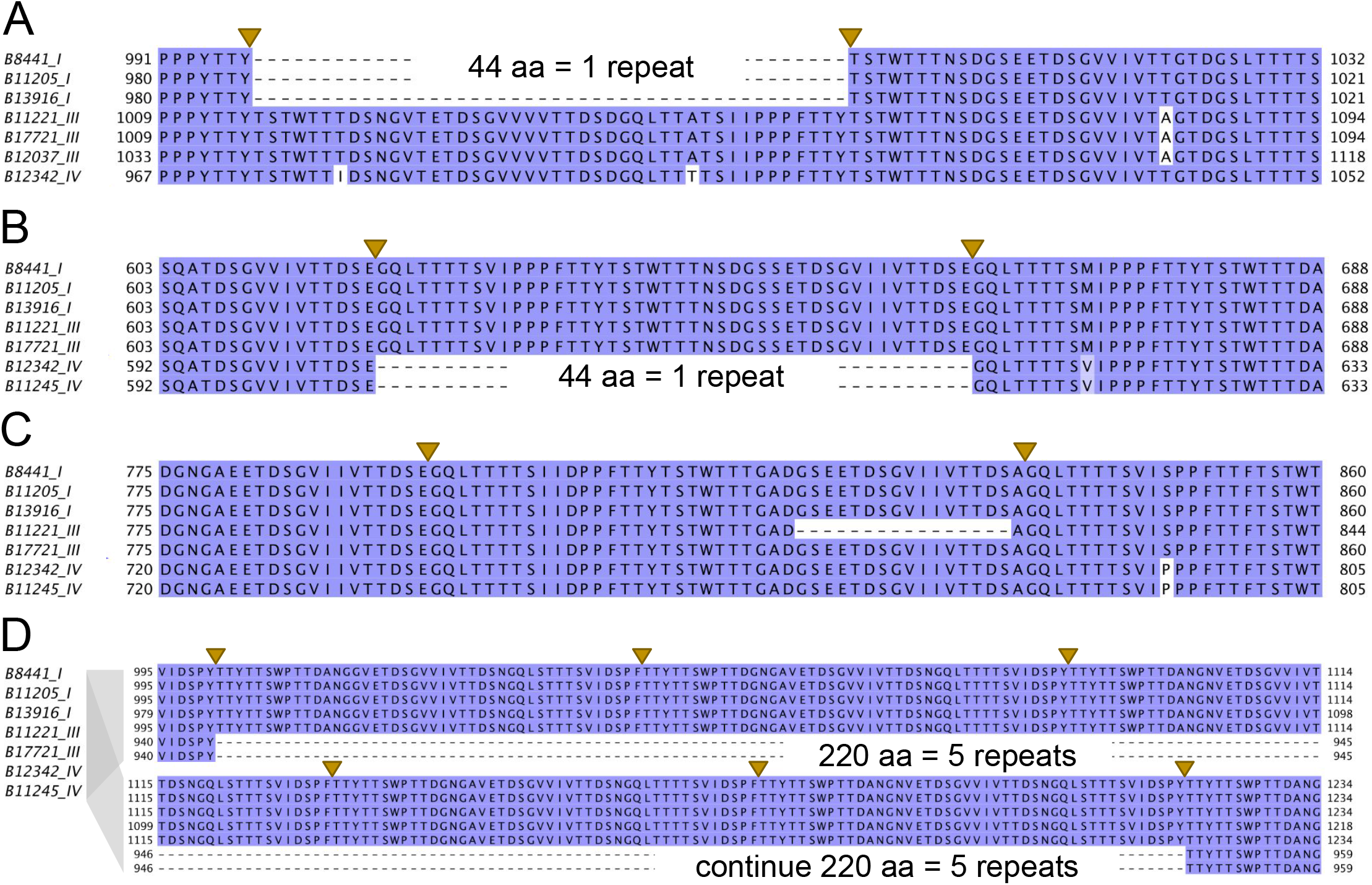
Examples of tandem repeat copy number variation in Hil1-Hil4 among the *C. auris* strains. (A) A 44 aa indel in Hil1 removes exactly one repeat in all three Clade I strain orthologs. (B) A similar indel polymorphism of exactly one repeat length in Hil2 affecting the Clade IV strains. (C) An indel polymorphism in Hil2 that affects one Clade III strain and spans 16 aa, not a full repeat, but includes a predicted strong β-aggregation prone sequence “GVIIVTT”. (D) An indel polymorphism in Hil2 that spans 220 aa or five full repeats affecting the Clade IV strains. Similar patterns were observed in Hil3 and Hil4.

**Supplementary Figure 6.**
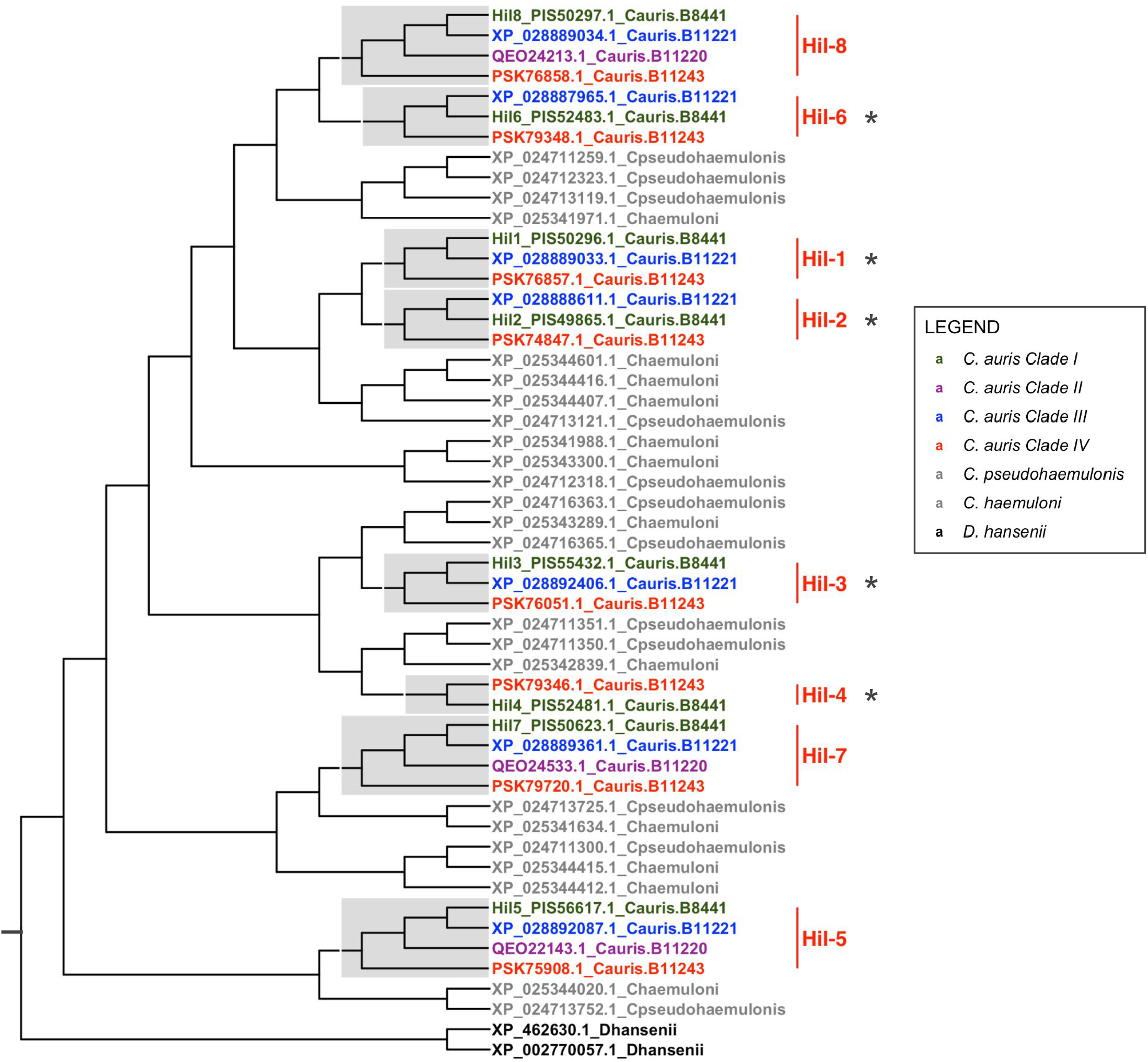
Reconciled Hil family gene tree based on the Hyphal_reg_CWP domain alignment in the four clades of *C. auris* strains and two closely related species. The tree is rooted by the two homologs from the outgroup *D. hansenii*. The gene tree was corrected with the species/strain tree based on (Muñoz et al 2018) using GeneRax (v2.0.4). Hil genes lost in *C. auris* Clade II strains are labeled with an asterisk next to the Hil1-8 group labels.

**Supplementary Figure 7.**
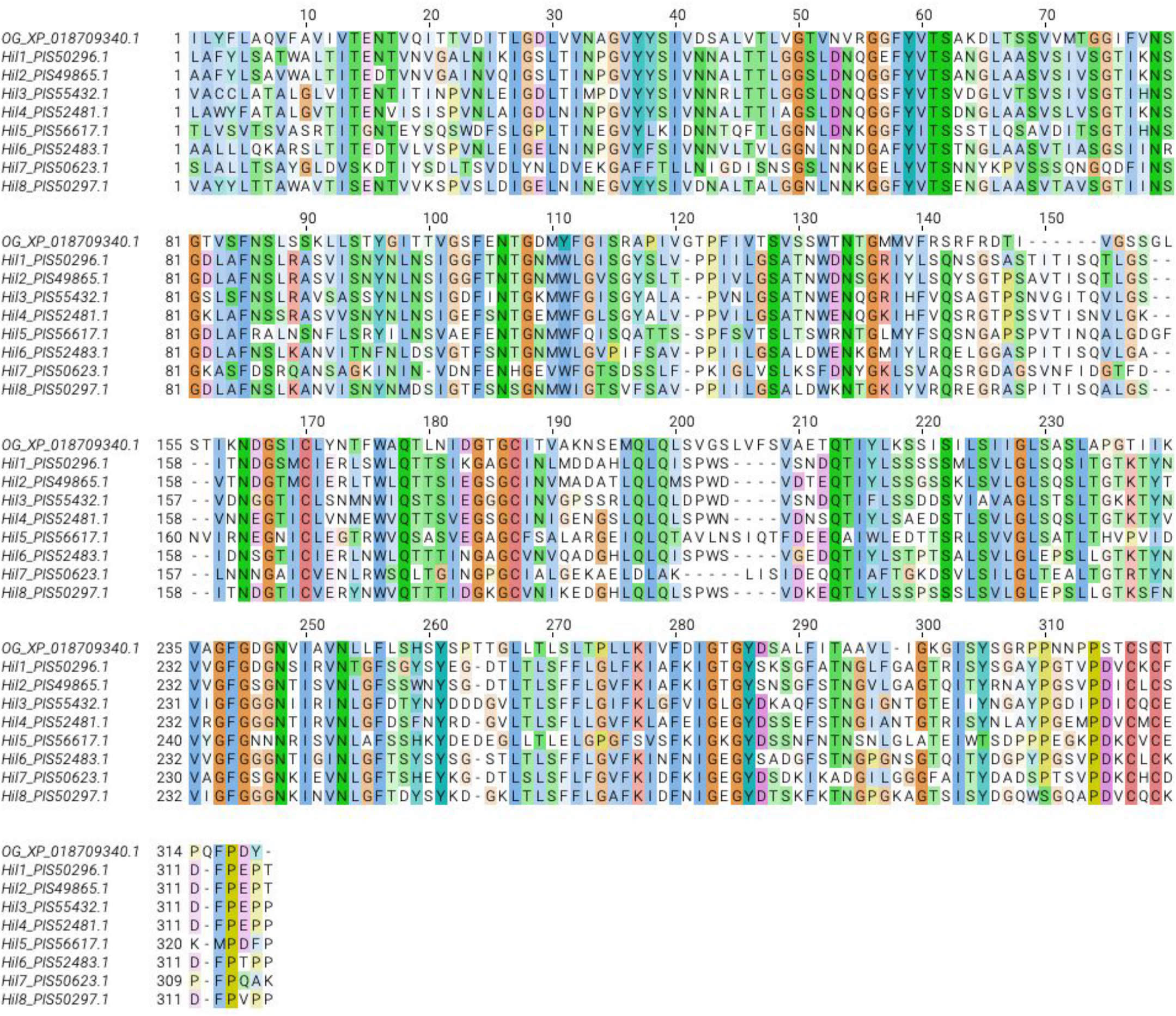
Multiple sequence alignment of *C. auris* Hil1-8 Hyphal_reg_CWP domain. The domain regions were identified using HMMScan against the Pfam-A database. The sequences were aligned with clustalo v1.2.4 and the result visualized in Jalview v2.11.1.4 with the ClustalW color scheme. OG = outgroup from *M. bicuspidata*

**Supplementary Figure 8.**
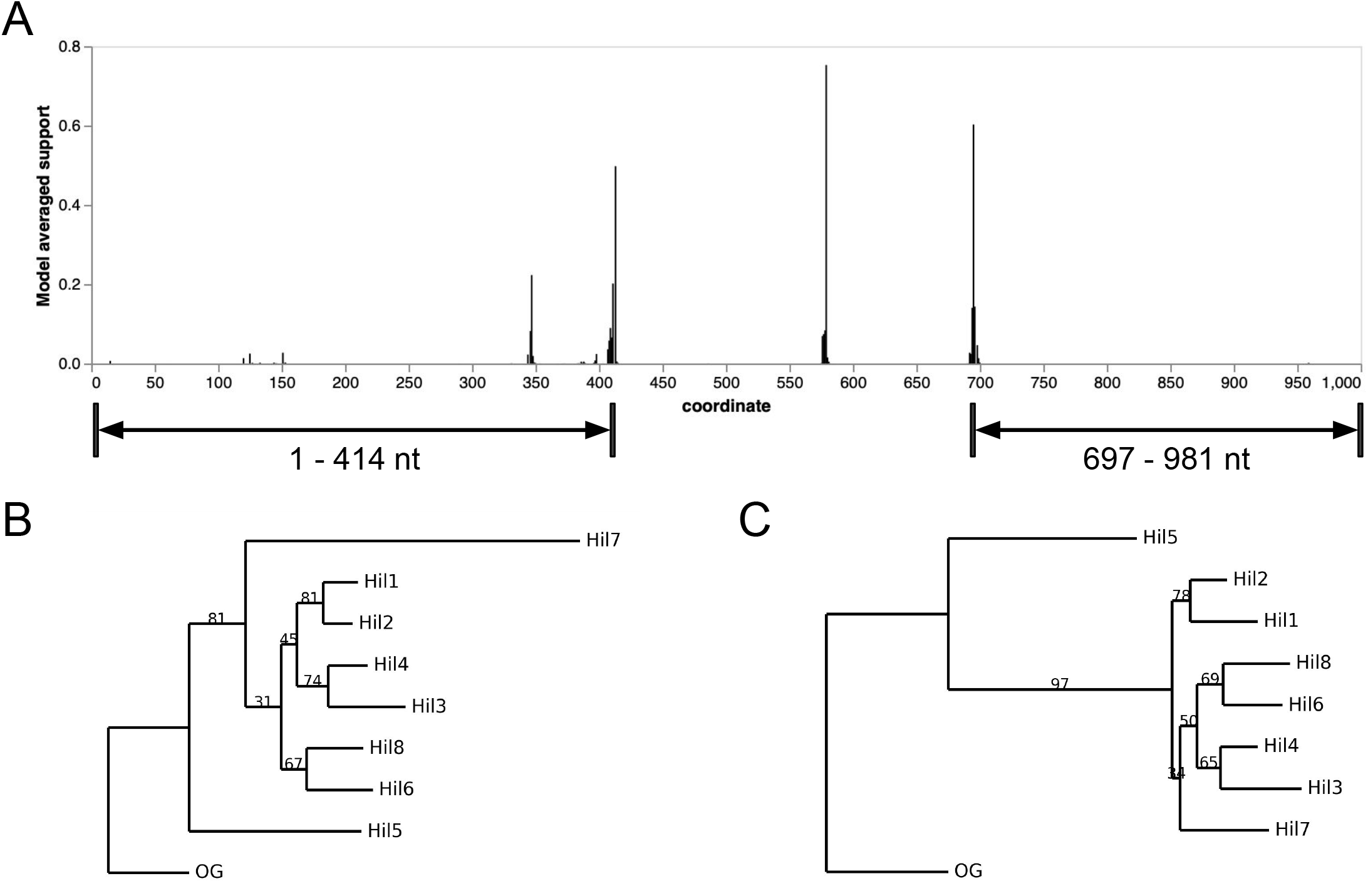
Detecting intra-domain recombination and identifying non-recombining partitions in the Hyphal_reg_CWP domain using GARD. (A) Model averaged support for breakpoint locations along the Hyphal_reg_CWP domain alignment for the eight Hil proteins in *C. auris* and an outgroup sequence from *M. bicuspidata* (protein ID: XP_018709340.1) to root the gene tree. Based on the GARD output, we chose the N- and C-terminal partitions for downstream analyses, i.e., coordinates 1-414 nt and 697-981 nt. (B) A maximum likelihood tree for partition 1-414 was constructed using RAxML-NG v1.1.0. Branch length is proportional to the amount of sequence divergence. OG stands for outgroup. Bootstrap support for internal splits are shown as a percentage and are based on 1000 replicates or until bootstrapping converges. (C) tree for 697-981nt, same format as in (B)

**Supplementary Figure 9.**
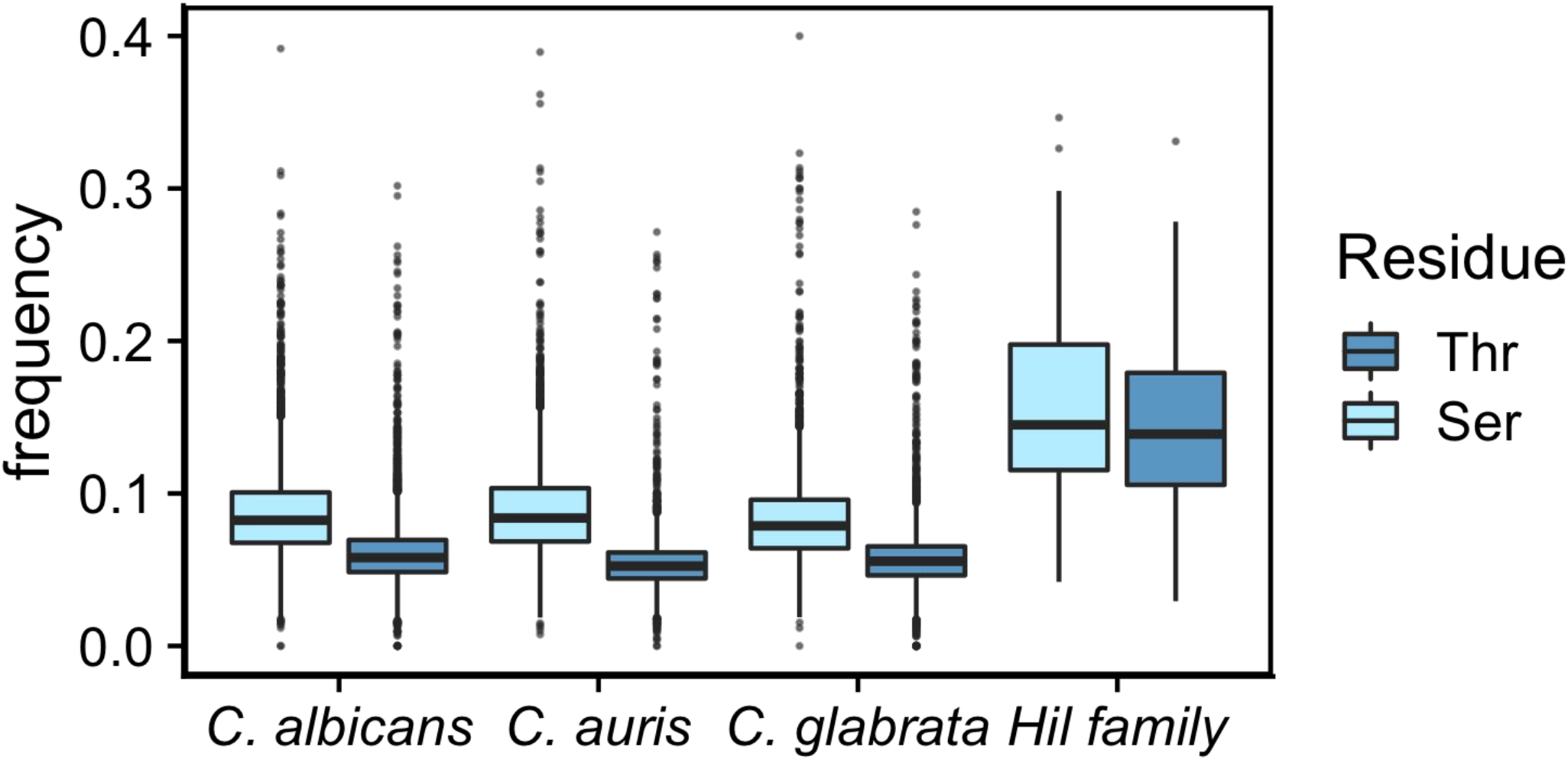
Yeast Hil family proteins have on average higher Ser/Thr frequencies than the rest of the proteome. Proteome-wide distribution of Thr/Ser frequencies per protein from three species, compared with the yeast Hil family proteins (*M. bicuspidata* homologs were excluded because a large number of them were incomplete). The boxes represent the interquartile range (IQR), the middle thick line the median, the whiskers the 1.5 x IQR and the dots outliers outside that range.

**Supplementary Figure 10.**
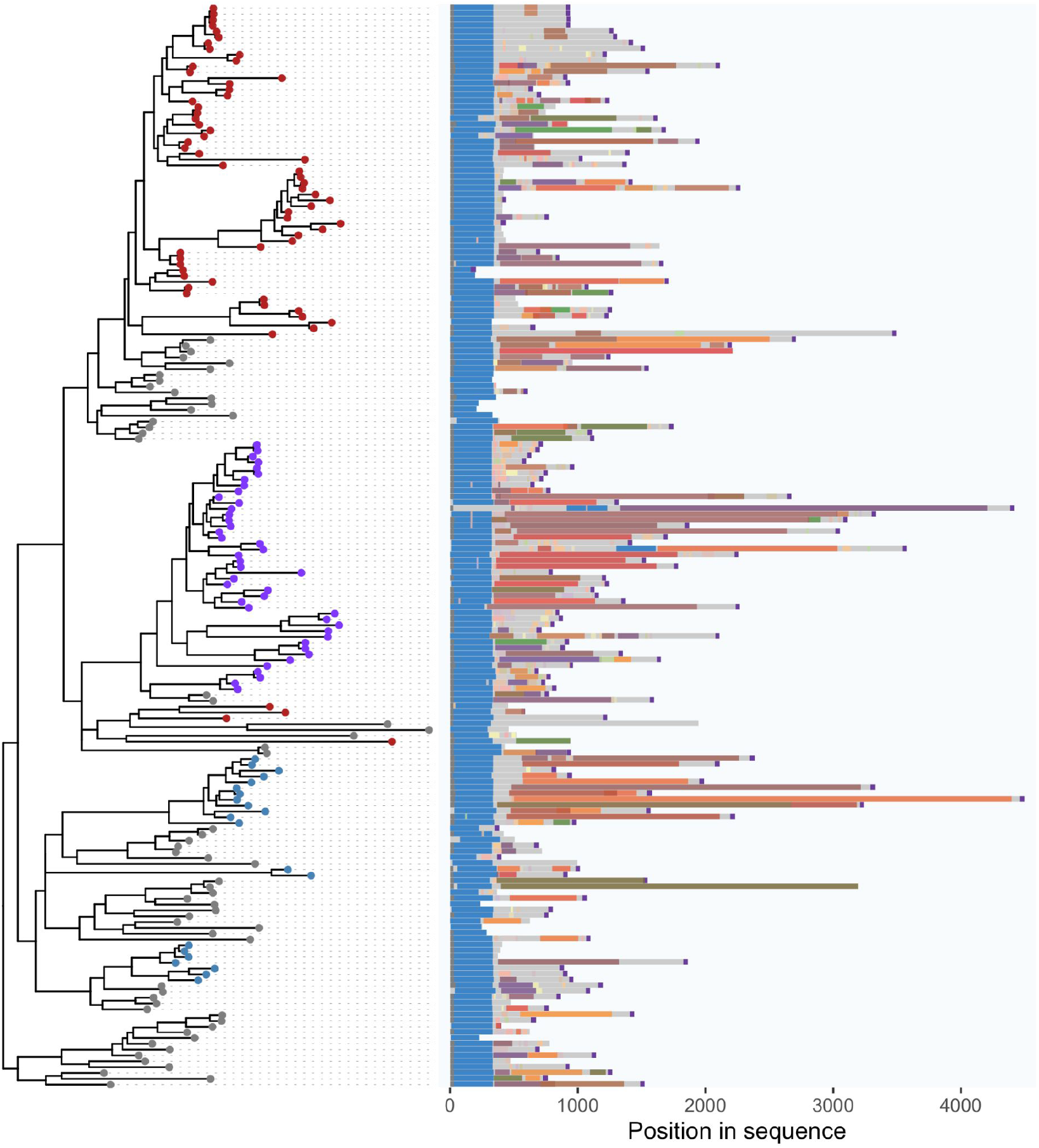
Domain schematic for the Yeast Hil family showing rapidly evolving tandem repeat sequences in the central domain of the proteins. Same as Fig. 6A except that tandem repeats belonging to different sequence clusters as determined by XSTREAM (Newman and Cooper 2007) are shown in different colors.

