## Supplementary material for "Parallel Expansion and Divergence of an Adhesin Family in Pathogenic Yeasts Including *Candida auris*": Text S1

### Text S1 - Identify Hil family homologs

Comprehensively identifying Hil family homologs and ensuring their accuracy are the basis for all downstream analyses. Therefore, we detail the multiple steps we took to minimize biases and errors. Despite all these efforts, we realize and note that our conclusions are only as good as the quality of the genome assemblies, which based on our analyses, are variable across species and in many cases need further improvement. Using new, long-read technologies such as Oxford Nanopore and PacBio in combination with Illumina sequences show great promise. But that alone is not sufficient. Careful bioinformatic processing, including assembly, manual curation and annotation, are all critical to accurate genome assemblies.

#### First round of identification using *C. auris* Hil1’s PF11765 domain as the query

To identify all Hil family homologs, we first used *C. auris* Hil1’s Hyphal_reg_CWP domain as the query and searched against the RefSeq database (as of May 2022) using BLASTP. Of the 189 hits that passed the *E*-value threshold of $10^{-5}$ and have a query coverage greater than 50%, all are from Ascomycota (yeasts) and all but one were from the Saccharomycetes class (budding yeast). A single hit was found in the fission yeast *Schizosacchromyces cryophilus*. Using that hit as the query, we searched all fission yeasts in the nr protein database with a relaxed E-value cutoff of $10^{-3}$, which resulted in no additional hit other than the query itself. Given this result, we conclude that the Hil family, defined by the presence of the Hyphal_reg_CWP domain, is yeast specific.

#### Expanded search using multiple Hil homologs’ PF11765 domain as queries

We next expanded the search with additional queries from two Hil family homologs, one from *C. albicans* Hyr1 and another from *C. glabrata* XP_445977. Based on an initial gene tree for all homologs identified in the first round, these three queries belong to different clades and span the entire tree. We reason that adding the *C. albicans* and *C. glabrata* queries could avoid biasing the homology identification to homologs more closely related to the *C. auris* query. The combined hit list with the same thresholds identified just three new hits compared with the first round, suggesting that our initial search included most of the homologous proteins.

#### Including additional yeast genome resources

The Genome Resources for Yeast Chromosomes (GRYC, http://gryc.inra.fr/) included additional yeast genomes not present in the NCBI RefSeq database, e.g., the Nakaseomyces genus that includes *Candida* pathogens closely related to *C. glabrata*. We performed the same search in this resource and recovered 16 additional homologs belonging to seven species.

#### Excluding species from the analysis

In order to infer the evolutionary history of the Hil family, including the duplication events, it is important to have a reliable species phylogeny. We relied on the recently published phylogeny for the budding yeast subphylum, which included 332 species (Shen et al. 2018). We manually added three species closely related to our focal species *C. auris*, i.e., *C. duobushaemulonis*, *C. pseudohaemulonii* and *C. haemuloni*, based on a separate phylogeny estimated by (Muñoz et al. 2018). By comparing the species included in the combined list of homologs from above, we excluded three species not in either species phylogeny, namely *Diutina rugosa*, *Kazachstania* *barnettii*, *Artibeus jamaicensis*. The remaining 32 species represent a broad sampling of the budding yeast subphylum. The final list of Hil family homologs includes 215 genes.

##### Verify RefSeq protein sequence annotation using newer assemblies

The RefSeq database provides a set of non-redundant and well-annotated sequences that can serve as a stable reference for gene identification and characterization. Because it emphasizes stability, newly sequenced and assembled genomes using more advanced technologies often take time to be integrated into the database. As a result, some of the RefSeq sequences may contain errors in their annotation. At a reviewer's suggestion, we performed TBLASTN search with the RefSeq hits in *C. tropicalis* against a long-read based assembly for the same MYA-3404 strain (assembly ID: GCA_013177555.1). We found several homologs had frameshift indels that postponed the stop codon further downstream in the new assembly. In those cases, we used the ORF finder (https://www.ncbi.nlm.nih.gov/orffinder/) to identify the longest ORF. Of the 16 homologs identified in this species, six were substantially longer in the new assembly than in the RefSeq assembly, and two were predicted to be substantially shorter (Table S8). Notably, while the new assembly is likely to be generally more reliable especially in the repeat-rich region of the Hil family homologs, sequencing and assembly errors are inevitably present in it as well. We then repeated the same strategy for other species that have a newer, long-read based assembly available, although not for the same strain as the RefSeq assembly, including *S. stipitis* (NRRL Y-7124, GCA_016859295.1), *C. albicans* (NCYC4166, GCA_005890745.1), *C. parapsilosis* (CBS6318, GCA_00982555.2) and *C. glabrata* (BG2, GCA_014217725.1). For *S. stipitis*, 16/18 RefSeq hits were shorter than 600 aa, and 15 were labeled as "incomplete". We identified seven hits with a query coverage above 50% in the new assembly, all of which are longer than 900 a.a. For the other three, the RefSeq hits are highly consistent in sequence with the new assembly, despite biological differences between strains. We conclude that the issues with the *C. tropicalis* and *S. stipitis* RefSeq hits are specific to their assemblies and not a general issue with the RefSeq database. More importantly, the sequence of the hits' PF11765 domain are always highly consistent between the RefSeq and the new assemblies, even for *C. tropicalis*. As our phylogenetic reconstruction is solely based on the alignment of the PF11765 domain, this means any assembly quality issues *do not* have an impact on the inference of the evolutionary history for the Hil family. Nonetheless, efforts to improve the genome assembly and update the RefSeq database are crucial for future studies to characterize gene family evolution, particularly when the gene family is repeat-rich.

##### Identify additional Hil family homologs in *C. glabrata* using the PacBio assembly

The Cormack lab recently sequenced and assembled the *C. glabrata* reference strains using PacBio (Xu et al. 2020), showing that the telomere and subtelomeric regions in the RefSeq assembly (GCF_000002545.3) were incomplete. To determine if our initial homology search missed any Hil family homologs in *C. glabrata*, we repeated the search in the new assembly (GCA_010111755.1). We identified a total of 13 hits passing all the criteria, three of which were identified in the RefSeq assembly. 12/13 hits are located in the subtelomeric regions, including all 10 that were identified in the new assembly alone. Since many yeast adhesin families, including the Hil family, tend to be located in subtelomeric regions, we repeated the homology search for additional species with a long-read assembly available, to determine how widespread the issue of missing homologs as observed in *C. glabrata* is. We focused on the genomes assembled at least to a chromosomal level, including *S. cerevisiae* (GCA_016858165.1), *K. lactis* (GCA_007993695.1), *C. nivariensis* (GCA_017309295.1) and *C. albicans* (GCA_005890745.1). In the first two species, the same number of hits (0 and 1) were identified in the long-read-based assembly as in the RefSeq one. In *C. nivariensis*, we identified three hits compared with two in the RefSeq assembly. Finally, in *C. albicans*, we identified 13 hits compared with 12 hits in the RefSeq assembly. However, two of the 13 hits were identical in the nucleotide sequence, raising questions as to whether they result from very recent duplications or assembly errors. This limited sample size suggests that the large discrepancy in the Hil family size observed in *C. glabrata* is unique and may have to do with the special challenges in assembling its subtelomeric regions. Nonetheless, we believe that improved assemblies for all species are urgently needed for studying adhesin families.
